# Structural basis of mechano-chemical coupling by the mitotic kinesin KIF14

**DOI:** 10.1101/2020.06.01.128371

**Authors:** Matthieu P.M.H. Benoit, Ana B. Asenjo, Mohammadjavad Paydar, Sabin Dhakal, Benjamin H. Kwok, Hernando Sosa

## Abstract

KIF14 is a mitotic kinesin protein important for cytokinesis. Its overexpression is associated with a variety of cancers and mutations in KIF14 result in cerebral and renal development defects. Like other kinesins, KIF14 contains a highly conserved catalytic motor domain where the energy from ATP hydrolysis is converted to directed movement along microtubules. Although much is known regarding the molecular mechanism of kinesin motility, there is a lack of structural information of kinesin-microtubule interactions at sufficient resolution to unambiguously assess how conformational changes related to ATP hydrolysis, microtubule binding and translocation are coupled. Here we determined the near-atomic resolution cryo-electron microscopy structures of five different KIF14 constructs bound to microtubules in the presence of different nucleotide analogues mimicking distinct steps of the ATPase cycle. Eighteen independent structures together with supporting functional assays provide a comprehensive view of the kinesin conformational changes occurring with microtubule and nucleotide binding. Our data shows that: 1) microtubule binding induces opening of the KIF14 nucleotide binding pocket; 2) AMP-PNP and ADP-AlF_x_ induce closure of the nucleotide binding pocket in microtubule bound KIF14 and this conformational change is allosterically controlled by the neck-linker domain; 3) the neck-linker domain when undocked prevents the nucleotide-binding-pocket to fully close and dampens ATP hydrolysis; 4) fifteen neck-linker residues are required to assume the docked conformation; 5) the nucleotide analogue ADP-AlF_x_ adopts a distinct configuration in an open nucleotide-binding-pocket; 6) the neck-linker position controls the hydrolysis step rather than nucleotide binding in the KIF14 ATPase cycle; 7) the two motor domains of a KIF14 dimer adopt distinct conformations when simultaneously bound to the microtubule. These observations provide the structural basis for a coordinated chemo-mechanical kinesin plus end translocation model.

## Introduction

KIF14 is a microtubule-based motor protein with essential roles during cell division (Barr and Gruneberg, 2007; Carleton et al., 2006; Gruneberg et al., 2006). Its overexpression is associated with tumor progression and resistance to anti-cancer drugs in several cancers for which is considered an oncogene (Corson and Gallie, 2006; Corson et al., 2005; Lucanus and Yip, 2017; Schiewek et al., 2018; Thériault et al., 2012; Wang et al., 2016). Mutations in KIF14 are also associated with neural and kidney development defects (Moawia et al., 2017; Reilly et al., 2018). However, despite the importance of KIF14 its action mechanism as a kinesin motor is still largely unknown.

KIF14 belongs to the kinesin-3 family of motor proteins. Other members of this family include KIF1A and CeUNC-104 (Miki et al., 2005). Kinesin-3s, in general work as microtubule plus end directed motors with the ability to make long processive runs when forming dimers (Soppina et al., 2014). KIF14 is also reported to be a microtubule plus end directed motile kinesin and to protect microtubules against depolymerization (Arora et al., 2014). As all kinesins, KIF14 possesses a highly conserved catalytic motor or head domain that contains nucleotide and microtubule binding sites and where ATP hydrolysis is coupled to the generation of mechanical work. The KIF14 molecule is also similar to other microtubule plus end directed motile kinesins in having two motor domain joined by a coiled coil dimerization domain located C-terminal to the motor domain and a ~15 residues long peptide, the neck-linker, connecting the motor and coiled coil domains (Verhey et al., 2015). KIF14 also contains an N-terminal extension with a PRC1 (protein-regulating cytokinesis 1) binding domain (Arora et al., 2014; Gruneberg et al., 2006; Reilly et al., 2018).

Although much is known regarding the mechanism of action of motile kinesins, and in particular of its founding member kinesin-1, most structural information of kinesin-microtubule complexes is still of limited resolution. As a consequence, it is still not fully clear how conformational changes are coupled to distinct steps of the ATP hydrolysis cycle or how the two heads of a kinesin dimer may coordinate their activities. To address this issue, we report here the near-atomic resolution structures of eighteen KIF14-microtubule complexes corresponding to five distinct KIF14 constructs in four nucleotide conditions mimicking four key steps of the ATP hydrolysis cycle. The data reveal how microtubule binding alters the structure of the KIF14 motor domain; how changes in nucleotide species bound to the catalytic site are coupled to motility-related conformational changes and how the two motor domains in a KIF14 dimer coordinate their activities. Our structural data provide a comprehensive molecular understanding of how kinesin ATPase cycle is coordinated with plus-end directed movement along microtubules.

## Results

### Cryo-electron microscopy structures of KIF14 motor domain complexes

To elucidate conformational changes related to the microtubule-bound KIF14 motor domain ATPase cycle, and how these may be modulated by the neck-linker or a partner motor domain, we determined the cryo-electron microscopy (cryo-EM) structures of microtubule complexes of five distinct KIF14 constructs in four nucleotide conditions. Full length mouse KIF14 is 1674 amino acids long and the core motor domain resides between residues asn-391 and ala-733 (Fig. 1a). The five constructs used in this work comprise residues asn-391 to asp-772 (K772), asn-391 to lys-755 (K755), asn-391 to ala-748 (K748), asn-391 to asn-743 (K743) and asn-391 to leu-735 (K735). All constructs include the core motor domain but differ in how many residues of the neck-linker (arg-734 to leu-750) and the dimerization coiled coil domain CC1 (ile-751 to ala-764, residues with > 0.99 coiled coil score according to COILS (Lupas et al., 1991)) they include. The longer K772 construct includes a full neck-linker and the first coiled coil domain CC1. Construct K755 includes the neck-linker and part of the first coiled coil heptad repeat of the CC1 domain. Construct K748 includes 15 neck-linker residues and no part of the CC1 domain. Construct K743 includes 10 neck-linker residues. Construct K735 comprises the core motor domain and two residues of the neck linker. We verified that construct K772 behaves in solution as a dimer while the shorter constructs behave as monomers (Fig. 1b,c). Structural snapshots in four distinct steps of the ATPase cycle were obtained by adding or removing nucleotides from the experimental solution as follow: 1) ADP added (4 mM) to induce the ADP bound state; 2) Traces of ATP and ADP removed with apyrase to induce the apo state; 3) Non-hydrolysable ATP analogue AMP-PNP added (4 mM) to mimic the ATP bound state; 4) ADP and aluminum fluoride added (4 mM ADP, 2 mM AlCl_3_, and 10 mM KF) to mimic the ADP-Pi transition state. The four nucleotide conditions will be denoted through the text as ADP, apo, ANP and AAF respectively.

**Figure 1.**
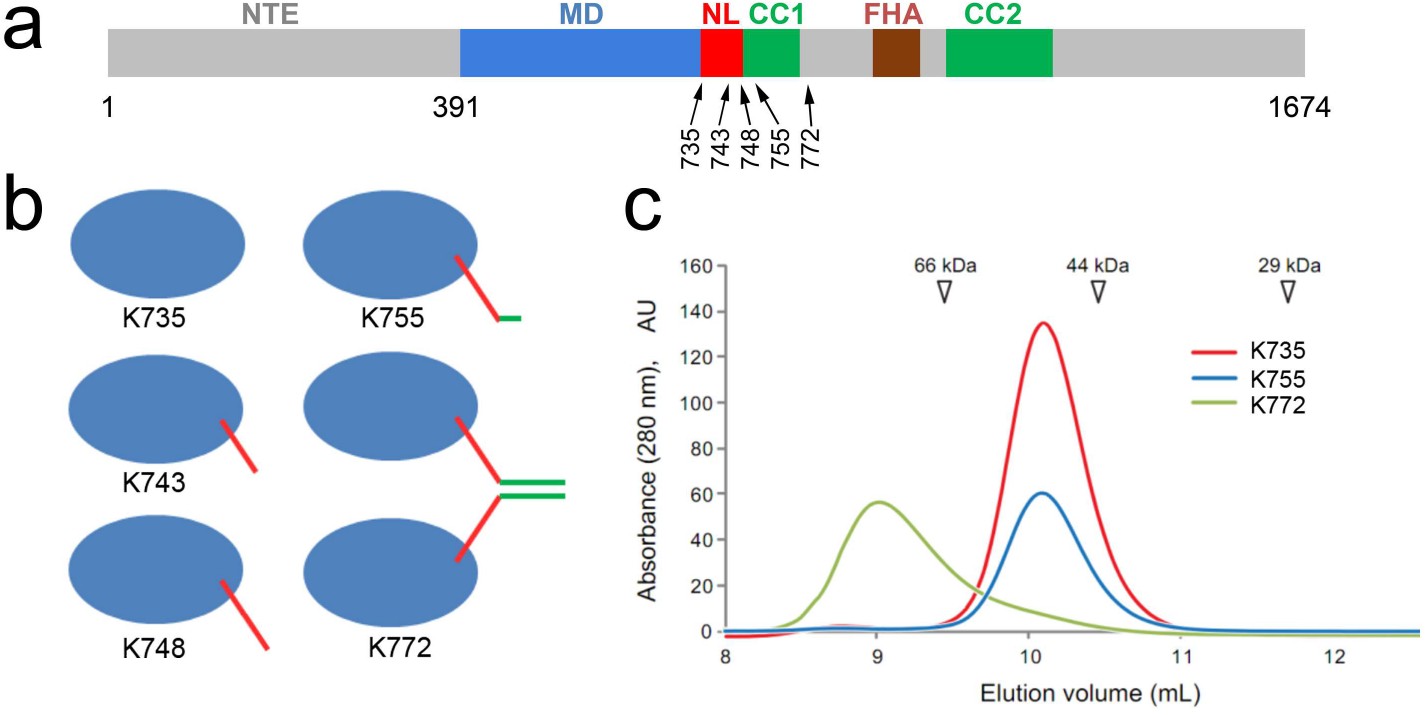
KIF14 constructs. **(a)** Primary structure and domain organization of *Mus musculu*s KIF14. NTE: N-terminal extension. MD: Motor domain. NL: Neck-linker domain. FHA: Forkhead-associated domain. CC1: Coiled coil domain 1. CC2: Coiled coil domain 2. Five constructs were used in this work, all include the motor domain from residue 391 at the N-terminal end and up to residues 735, 743, 748, 755 or 772 (relative size of bars and arrow positions not at scale). **(b)** Predicted quaternary structure of the KIF14 constructs used. **(c)** Gel filtration profile of constructs K735, K755, and K772.

Combining the different KIF14 constructs and nucleotide conditions we calculated the near-atomic cryo-EM structures of eighteen distinct microtubule-KIF14 complexes (Table 1, Fig. 2). All the KIF14-microtubule cryo-EM structures show the typical kinesin motor domain decoration pattern with one motor domain bound to each tubulin heterodimer in the microtubule lattice (Fig. 2). The attained resolution of the cryo-EM maps, in the 2.9-4.0 Å range (Table 1, Supplementary Fig. 1), allowed us to generate accurate atomic models for all the conditions investigated, trace the polypeptide chains, locate most amino-acid side chains and clearly identify the nucleotide species present in the nucleotide binding pocket. From the eighteen cryo-EM maps we generated twenty-two distinct atomic models of the KIF14-motor-domain-microtubule complex (Tables 2, 3). Four maps, MT-K755-ANP, MT-K755-AAF, MT-K772-ANP and MT-K772-AAF were fitted with two coexisting motor domain structures (Leading and Trailing heads). We found significant structural differences between the complexes related to the KIF14-microtubule interaction, the nucleotide species present and whether the construct includes the neck-linker or dimerization domains. The twenty-two structures together with the previously solved crystal structure of the microtubule unbound KIF14 motor domain (Arora et al., 2014) can be grouped into four groups based on their structural similarity (Tables 2, 3). In the following sections we describe and discuss the significance of all the KIF14 structures solved and their differences.

**Table 1.** Cryo-EM data collection refinement and validation statistics.

|  | K735-ANP<br>(EMDB-21949)<br>(PDB 6WWV) | K735-AAF<br>(EMDB-21948)<br>(PDB 6WWU) | K735 apo<br>(EMDB-21947)<br>(PDB 6WWT) | K743-ANP<br>(EMDB-21946)<br>(PDB 6WWS) |
| --- | --- | --- | --- | --- |
| **Data collection and processing** |  |  |  |  |
| Magnification (actual) | 46598 | 60168 | 46598 | 58893 |
| Voltage (kV) | 300 | 300 | 300 | 300 |
| Electron exposure (e-/Å<sup>2</sup>) | 69.9 | 70.1 | 69.9 | 63.0 |
| Defocus range (µm)<sup>a</sup> | 1.0 - 1.9 | 1.1 - 1.9 | 1.0 - 2.0 | 0.9 - 1.6 |
| Pixel size (Å) | 1.073 | 0.831 | 1.073 | 0.849 |
| Symmetry imposed | Helical | Helical | Helical | Helical |
| Rise (Å) | 5.5 | 5.41 | 5.46 | 5.45 |
| Twist (deg) | 168.09 | 168.07 | 168.09 | 168.09 |
| Particle images identified as 15R symmetry (no.) | 15350 | 18849 | 9863 | 38121 |
| Final particle images (no.) | 15350 | 18849 | 9863 | 38121 |
| Number of asymmetrical units | 228854 | 285696 | 148125 | 573563 |
| Map resolution (Å) | 3.5 | 3.0 | 3.6 | 3.0 |
| FSC threshold | 0.143 | 0.143 | 0.143 | 0.143 |
| Kinesin resolution (Å) | 3.7 | 3.2 | 3.8 | 3.2 |
| Tubulin resolution (Å) | 3.4 | 3.0 | 3.5 | 2.9 |
| **Refinement** |  |  |  |  |
| Initial model used (PDB code) | 6B0I | 6B0I | 6B0I | 6B0I |
| Model composition |  |  |  |  |
| Non-hydrogen atoms | 9677 | 9680 | 9645 | 9686 |
| Protein residues | 1215 | 1216 | 1215 | 1216 |
| Ligands | 4 | 5 | 3 | 4 |
| R.m.s. deviations |  |  |  |  |
| Bond lengths (Å) | 0.011 | 0.009 | 0.009 | 0.01 |
| Bond angles (°) | 1.32 | 1.18 | 1.16 | 1.20 |
| Validation |  |  |  |  |
| MolProbity score | 2.28 | 2.03 | 1.89 | 1.98 |
| Clashscore | 8.56 | 6.72 | 5.90 | 6.30 |
| Poor rotamers (%) | 2.68 | 2.01 | 1.63 | 1.82 |
| Ramachandran plot |  |  |  |  |
| Favored (%) | 91.73 | 93.55 | 93.96 | 93.31 |
| Allowed (%) | 7.61 | 6.45 | 5.79 | 6.53 |
| Disallowed (%) | 0.66 | 0 | 0.25 | 0.17 |

Notes:
<sup>a</sup> Range of the average defocus measured on the particle images used. The range comprises 90% of the particles used (5% of the particle defocuses are below and 5% are above this range).

Table 1 (3/5).
|  | K743-AAF<br>(EMDB-21945)<br>(PDB 6WWR) | K743-ADP<br>(EMDB-21944)<br>(PDB 6WWQ) | K743 apo<br>(EMDB-21943)<br>(PDB 6WWP) | K748-ANP<br>(EMDB-21942)<br>(PDB 6WWO) |
| --- | --- | --- | --- | --- |
| **Data collection and processing** |  |  |  |  |
| Magnification (actual) | 60606 | 60386 | 58962 | 60386 |
| Voltage (kV) | 300 | 300 | 300 | 300 |
| Electron exposure (e-/Å<sup>2</sup>) | 70.8 | 70.2 | 70.0 | 68.2 |
| Defocus range (µm) <sup>a</sup> | 0.8 – 1.7 | 0.8 – 1.6 | 0.9 – 1.8 | 0.9 – 1.7 |
| Pixel size (Å) | 0.825 | 0.828 | 0.848 | 0.828 |
| Symmetry imposed | Helical | Helical | Helical | Helical |
| Rise (Å) | 5.44 | 5.42 | 5.45 | 5.43 |
| Twist (deg) | 168.07 | 168.08 | 168.09 | 168.08 |
| Particle images identified as 15R symmetry (no.) | 41714 | 5194 | 12644 | 16440 |
| Final particle images (no.) | 41714 | 5194 | 12644 | 11075 |
| Number of asymmetrical units | 628777 | 78580 | 190240 | 167246 |
| Map resolution (Å) | 2.9 | 3.3 | 3.3 | 3.2 |
| FSC threshold | 0.143 | 0.143 | 0.143 | 0.143 |
| Kinesin resolution (Å) | 3.2 | 3.8 | 3.6 | 3.5 |
| Tubulin resolution (Å) | 2.9 | 3.3 | 3.2 | 3.1 |
| **Refinement** |  |  |  |  |
| Initial model used (PDB code) | 6B0I | 6B0I | 6B0I | 6B0I |
| Model composition |  |  |  |  |
| Non-hydrogen atoms | 9711 | 9675 | 9634 | 9784 |
| Protein residues | 1220 | 1215 | 1213 | 1230 |
| Ligands | 5 | 4 | 3 | 4 |
| R.m.s. deviations |  |  |  |  |
| Bond lengths (Å) | 0.011 | 0.011 | 0.008 | 0.011 |
| Bond angles (°) | 1.28 | 1.12 | 1.15 | 1.29 |
| Validation |  |  |  |  |
| MolProbity score | 2.34 | 1.89 | 2.24 | 2.13 |
| Clashscore | 8.43 | 7.78 | 6.75 | 7.01 |
| Poor rotamers (%) | 3.15 | 1.05 | 3.26 | 2.65 |
| Ramachandran plot |  |  |  |  |
| Favored (%) | 91.60 | 93.13 | 92.21 | 93.55 |
| Allowed (%) | 8.24 | 6.70 | 7.46 | 6.13 |
| Disallowed (%) | 0.16 | 0.17 | 0.33 | 0.33 |

Table 1 (4/5).
|  | K748-AAF<br>(EMDB-21941)<br>(PDB 6WWN) | K748-ADP<br>(EMDB-21940)<br>(PDB 6WWM) | K755 ANP<br>(EMDB-21939)<br>(PDB 6WWL) | K755-AAF<br>(EMDB-21938)<br>(PDB 6WWK) |
| --- | --- | --- | --- | --- |
| **Data collection and processing** |  |  |  |  |
| Magnification (actual) | 60606 | 58893 | 46624 | 58962 |
| Voltage (kV) | 300 | 300 | 300 | 300 |
| Electron exposure (e-/Å<sup>2</sup>) | 70.4 | 62.4 | 72.1 | 73.4 |
| Defocus range (µm) <sup>a</sup> | 1.0 - 1.7 | 0.8 - 1.9 | 0.6 - 2.0 | 0.6 - 1.7 |
| Pixel size (Å) | 0.825 | 0.849 | 1.0724 | 0.848 |
| Symmetry imposed | Helical | Helical | Helical | Helical |
| Rise (Å) | 5.41 | 5.51 | 5.47 | 5.45 |
| Twist (deg) | 168.07 | 168.09 | 168.09 | 168.07 |
| Particle images identified as 15R symmetry (no.) | 15178 | 14102 | 32006 | 26624 |
| Final particle images (no.) | 15178 | 14102 | 32006 | 26624 |
| Number of asymmetrical units | 230054 | 209866 | 479797 | 400581 |
| Map resolution (Å) | 3.9 | 3.1 | 3.4 | 3.2 |
| FSC threshold | 0.143 | 0.143 | 0.143 | 0.143 |
| Kinesin resolution (Å) | 3.9 | 3.5 | 3.5 | 3.5 |
| Tubulin resolution (Å) | 3.9 | 3.1 | 3.3 | 3.1 |
| **Refinement** |  |  |  |  |
| Initial model used (PDB code) | 6B0I | 6B0I | 6B0I | 6B0I |
| Model composition |  |  |  |  |
| Non-hydrogen atoms | 9784 | 9661 | 19666 | 19690 |
| Protein residues | 1230 | 1213 | 2470 | 2473 |
| Ligands | 5 | 4 | 8 | 10 |
| R.m.s. deviations |  |  |  |  |
| Bond lengths (Å) | 0.009 | 0.009 | 0.010 | 0.009 |
| Bond angles (°) | 1.21 | 1.14 | 1.1 | 1.16 |
| Validation |  |  |  |  |
| MolProbity score | 2.02 | 1.72 | 2.08 | 2.19 |
| Clashscore | 6.34 | 4.89 | 5.91 | 5.73 |
| Poor rotamers (%) | 1.90 | 1.06 | 2.45 | 4.61 |
| Ramachandran plot |  |  |  |  |
| Favored (%) | 92.81 | 92.96 | 92.64 | 94.63 |
| Allowed (%) | 7.11 | 6.96 | 7.32 | 5.33 |
| Disallowed (%) | 0.08 | 0.08 | 0.04 | 0.04 |

Table 1 (5/5).
|  | K755-ADP<br>(EMDB-21937)<br>(PDB 6WWJ) | K755 apo<br>(EMDB-21936)<br>(PDB 6WWI) | K772ANP<br>(EMDB-21935)<br>(PDB 6WWH) | K772-AAF<br>(EMDB-21934)<br>(PDB 6WWG) |
| --- | --- | --- | --- | --- |
| **Data collection and processing** |  |  |  |  |
| Magnification (actual) | 60168 | 45956 | 46598 | 60533 |
| Voltage (kV) | 300 | 300 | 300 | 300 |
| Electron exposure (e-/Å<sup>2</sup>) | 70.6 | 69.2 | 67.3 | 75.1 |
| Defocus range (µm) <sup>a</sup> | 0.7 - 1.7 | 0.7 - 1.8 | 0.8 - 2.0 | 0.7 - 1.6 |
| Pixel size (Å) | 0.831 | 1.088 | 1.073 | 0.826 |
| Symmetry imposed | Helical | Helical | Helical | Helical |
| Rise (Å) | 5.46 | 5.5 | 5.48 | 5.44 |
| Twist (deg) | 168.09 | 169.09 | 168.09 | 168.07 |
| Particle images identified as 15R symmetry (no.) | 18744 | 15798 | 3395 | 23479 |
| Final particle images (no.) | 18744 | 15798 | 3395 | 23479 |
| Number of asymmetrical units | 281503 | 235533 | 50801 | 353911 |
| Map resolution (Å) | 3.5 | 3.8 | 4.0 | 3.0 |
| FSC threshold | 0.143 | 0.143 | 0.143 | 0.143 |
| Kinesin resolution (Å) | 3.9 | 4.1 | 4.3 | 3.3 |
| Tubulin resolution (Å) | 3.5 | 3.8 | 3.9 | 3.0 |
| **Refinement** |  |  |  |  |
| Initial model used (PDB code) | 6B0I | 6B0I | 6B0I | 6B0I |
| Model composition |  |  |  |  |
| Non-hydrogen atoms | 9786 | 9753 | 19685 | 19773 |
| Protein residues | 1230 | 1229 | 2471 | 2484 |
| Ligands | 4 | 3 | 8 | 10 |
| R.m.s. deviations |  |  |  |  |
| Bond lengths (Å) | 0.011 | 0.013 | 0.009 | 0.011 |
| Bond angles (°) | 1.16 | 1.07 | 1.12 | 1.12 |
| Validation |  |  |  |  |
| MolProbity score | 1.71 | 1.93 | 2.09 | 2.00 |
| Clashscore | 4.62 | 7.18 | 6.60 | 5.19 |
| Poor rotamers (%) | 1.33 | 1.04 | 2.64 | 2.21 |
| Ramachandran plot |  |  |  |  |
| Favored (%) | 94.44 | 91.01 | 94.02 | 92.64 |
| Allowed (%) | 5.31 | 8.99 | 5.82 | 7.24 |
| Disallowed (%) | 0.25 | 0 | 0.16 | 0.12 |

**Table 1. Cryo-EM data collection refinement and validation statistics.**
|  | K772-ADP<br>(EMDB-21933)<br>(PDB 6WWF) | K772 apo<br>(EMDB-21932)<br>(PDB 6WWE) |
| --- | --- | --- |
| **Data collection and processing** |  |  |
| Magnification (actual) | 58893 | 58824 |
| Voltage (kV) | 300 | 300 |
| Electron exposure (e-/Å<sup>2</sup>) | 66.8 | 68.7 |
| Defocus range (µm)<sup>a</sup> | 0.5 - 1.5 | 0.7 - 1.9 |
| Pixel size (Å) | 0.849 | 0.850 |
| Symmetry imposed | Helical | Helical |
| Rise (Å) | 5.48 | 5.50 |
| Twist (deg) | 168.09 | 168.08 |
| Particle images identified as 15R symmetry (no.) | 7377 | 3843 |
| Final particle images (no.) | 7377 | 3843 |
| Number of asymmetrical units | 110385 | 21008 |
| Map resolution (Å) | 3.5 | 3.9 |
| FSC threshold | 0.143 | 0.143 |
| Kinesin resolution (Å) | 3.8 | 4.4 |
| Tubulin resolution (Å) | 3.4 | 3.8 |
| **Refinement** |  |  |
| Initial model used (PDB code) | 6B0I | 6B0I |
| Model composition |  |  |
| Non-hydrogen atoms | 9793 | 9765 |
| Protein residues | 1231 | 1231 |
| Ligands | 4 | 3 |
| R.m.s. deviations |  |  |
| Bond lengths (Å) | 0.010 | 0.01 |
| Bond angles (°) | 1.07 | 1.09 |
| Validation |  |  |
| MolProbity score | 1.84 | 2.09 |
| Clashscore | 6.43 | 10.04 |
| Poor rotamers (%) | 0.38 | 0.09 |
| Ramachandran plot |  |  |
| Favored (%) | 91.92 | 89.39 |
| Allowed (%) | 8.08 | 10.45 |
| Disallowed (%) | 0 | 0.16 |

**Figure 2.**
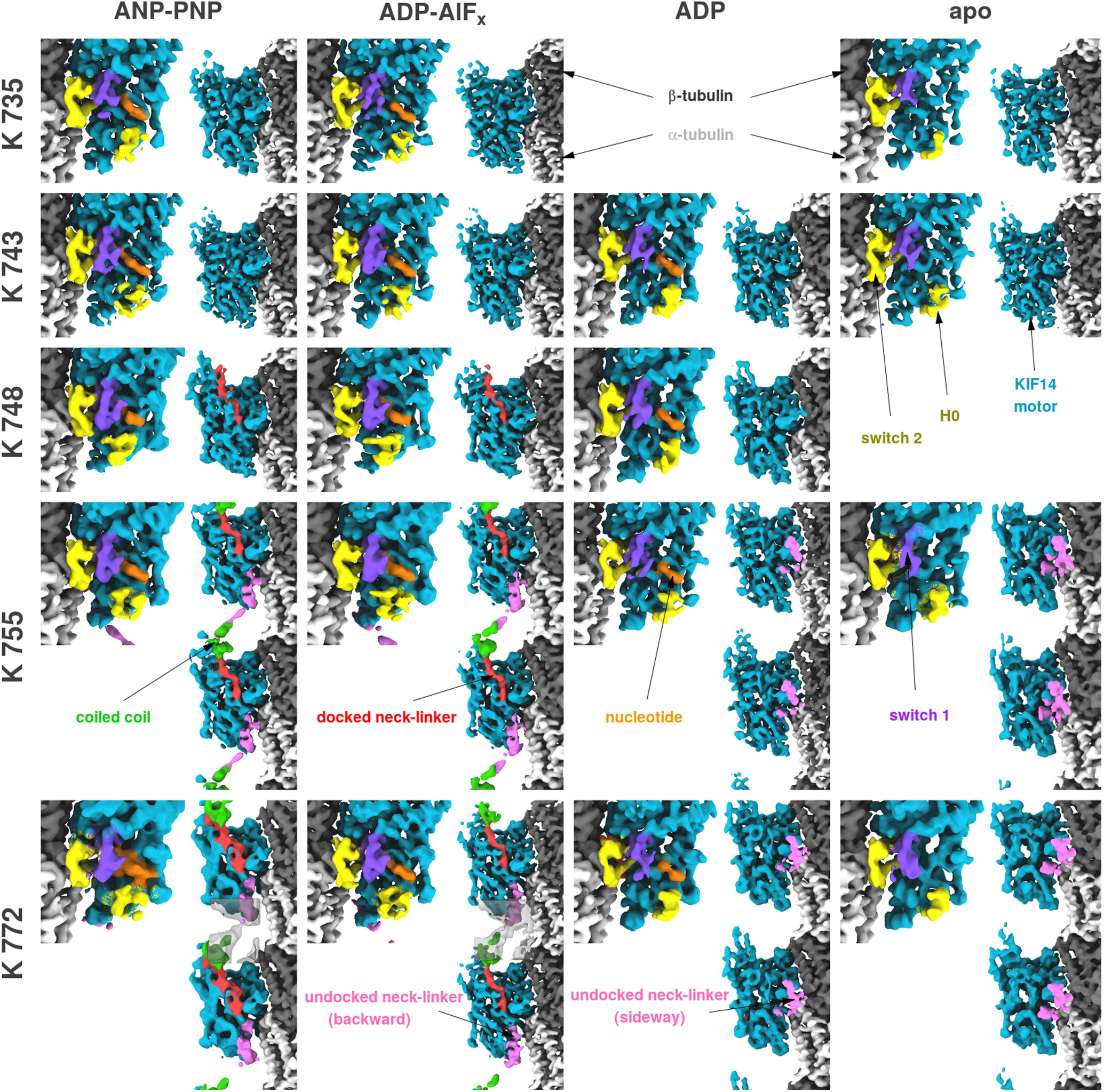
Cryo-EM 3D density maps. Iso-density surface representations of the eighteen-microtubule bound KIF14 complexes cryo-EM maps determined. The columns correspond to the four different nucleotide conditions used and the rows to the five KIF14 constructs used. For each construct/nucleotide combination the left panel show a zoom of the nucleotide binding pocket of the kinesin and the right panel shows the neck linker area at lower magnification. Both views are rotated 180 degree from each other. Surfaces are colored according to the fitted atomic structure they enclose: α-tubulin in light gray, β-tubulin in dark gray, most of the kinesin motor domain in blue, the switch-1 loop in purple, the swtch-2 loop in yellow, KIF14 α-helix-0 in yellow, the bound nucleotide (ADP or AMP-PNP) in orange, the neck-linker when docked in red and when undocked in pink and the CC1 coiled coil in green. Densities in the undocked neck linker (pink) and coiled coil (green) are only present in the K755 and K772 constructs but are noisier than in the rest of the KIF14 motor domain area and were displayed at a lower contour level. For the maps with the K772 construct with ANP-PNP and ADP-AlF_x_ a low-pass filtered version (semi-transparent grey surface) was superimposed to better display the noisier densities connecting the motor domains. The figure was prepared with VMD (Humphrey et al., 1996).

**Table 2.**
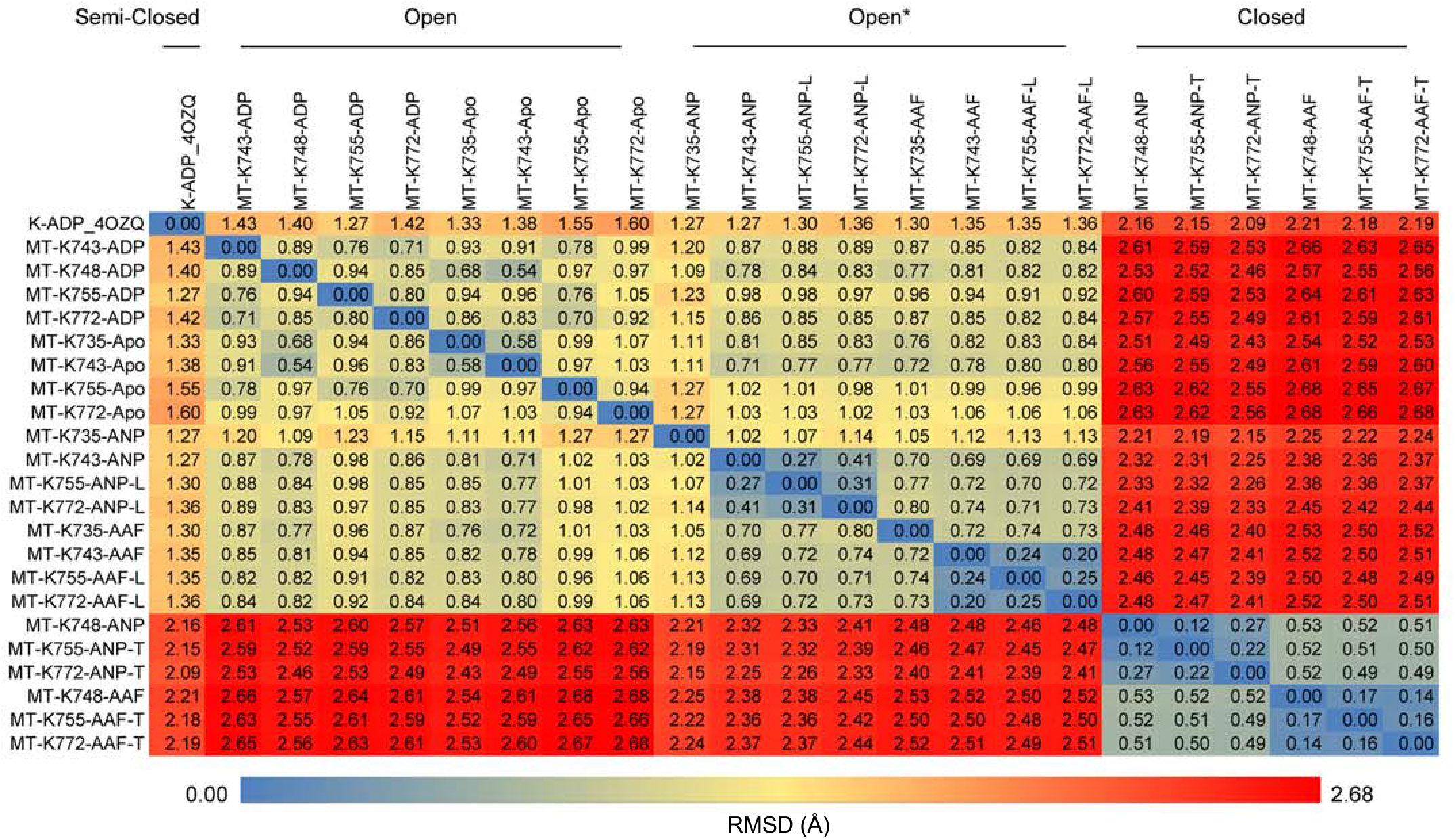
KIF14 motor domain structures pairwise comparison. The table shows the root mean square distance (RMSD) in Å, after optimal alignment of equivalent C-α carbons of the KIF14 motor domain of the two complexes specified in the corresponding row and column. Only residues resolved in all the structures were used in the alignment and RMSD calculations. This excludes part of the tubulin interacting loops and the tips of the switch loops not visible in the microtubule-unbound crystal structure, as well as the neck-linker which is only visible in some microtubule bound-structures. Table cells are color coded according to their RMSD values.

**Table 3.**
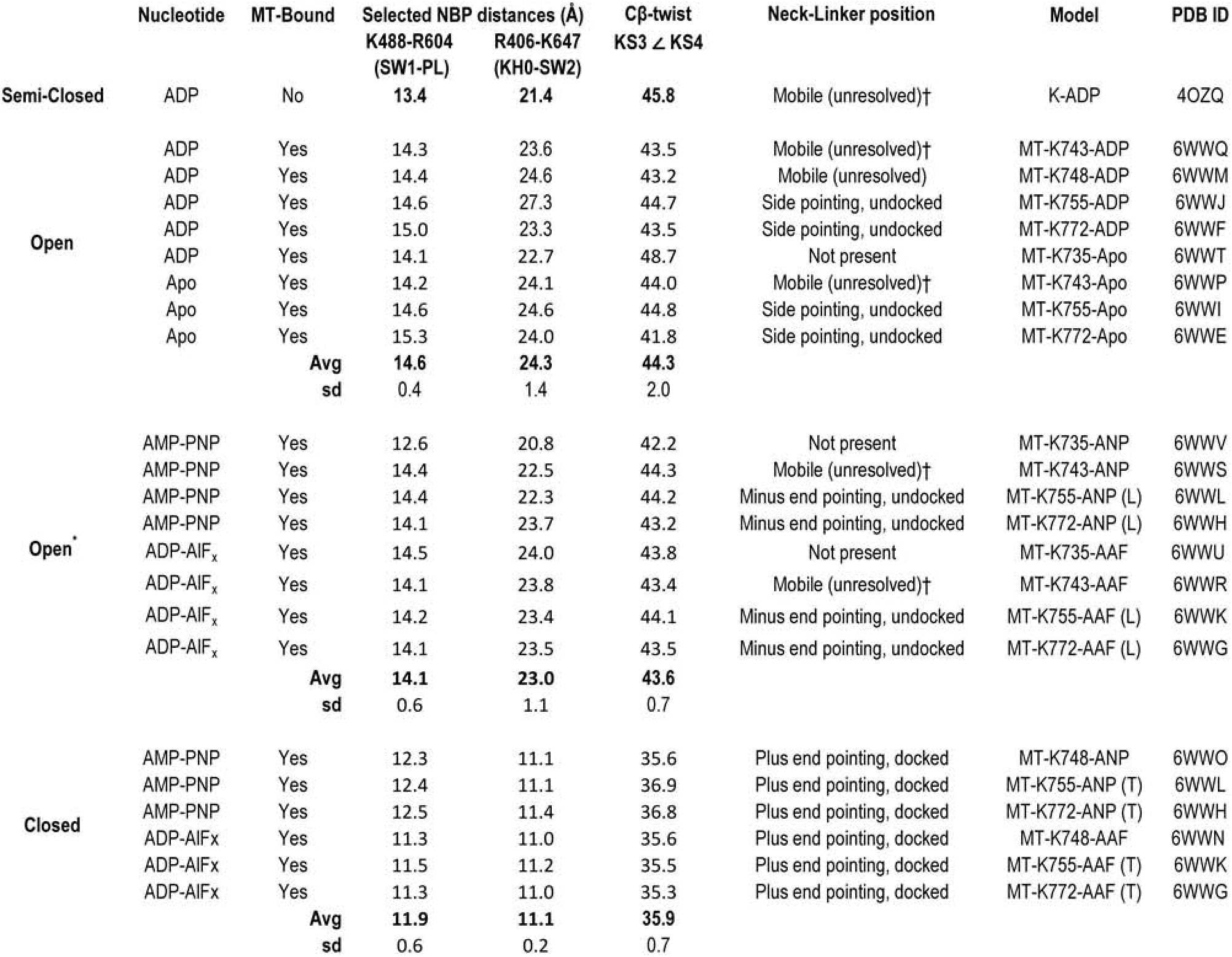
KIF14 conformation groups key features. Selected nucleotide-binding-pocket (NBP) distances correspond to the distance between Cα carbons in selected residues around the NBP. Central β-sheet twist (Cβ-twist) was measured as the angle between two vectors formed by the coordinates of the Cα carbons of KIF14 residues 475 and 480 in KS3 and residues 527 and 531 in KS4 (Fig. 3f). ^†^ Indicates constructs with truncated neck-linkers. Construct K743 has the neck-linker truncated at position 743 and the microtubule unbound KIF14 crystal structure (PDB: 4OZQ) at residue 738.

### Microtubule binding opens the KIF14 nucleotide binding pocket enabling nucleotide exchange

To examine the effect of microtubule binding on the structure of the KIF14 motor domain, we compared the microtubule-bound cryo-EM structures with the crystal structure of the ADP bound KIF14 motor domain (Arora et al., 2014). The first thing we noticed is that the interaction with the microtubule induces order in regions of the motor domain near the interface with the microtubule. In the microtubule unbound KIF14 crystal structure, parts of the kinesin loops 8, 9 and 11 (KL8, KL9, KL11) are not resolved and thus presumed disordered. On the other hand, the microtubule bound cryo-EM structures show clear densities along the full length of these regions (Fig. 3a). KL9 and KL11 correspond to the switch loops (SW1 and SW2). These are common structural elements in the related NTPases, kinesins, myosins and G-proteins, that sense the nucleotide species in the active site (Vale, 1996).

**Figure 3.**
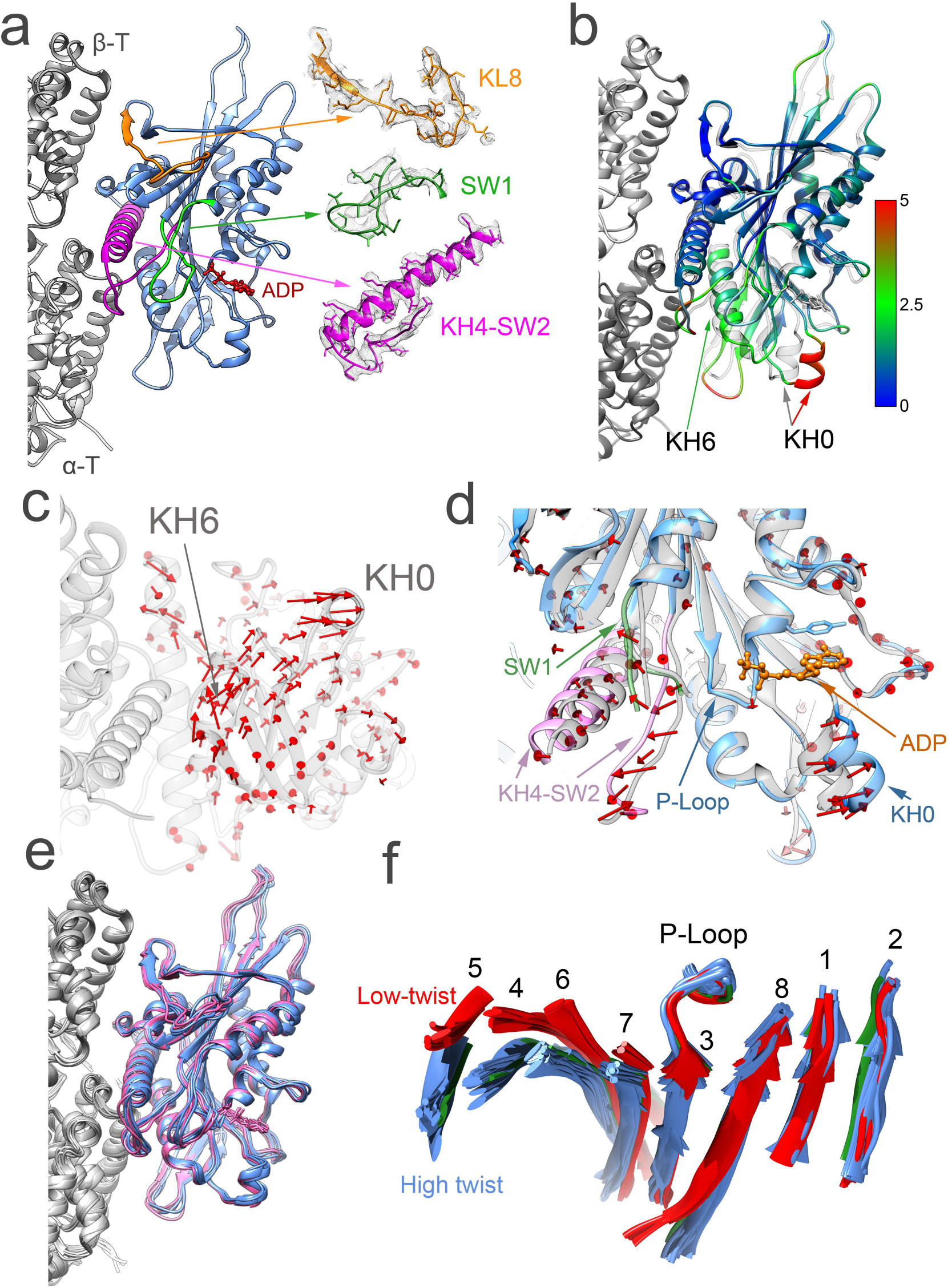
Microtubule bound ADP and apo structures. **(a)** Side view of MT-K748-ADP atomic model structure. Insets show regions at the KIF14-microtubule interface (KL8) and the switch loops (SW1 and SW2) with corresponding experimental density (gray mesh). **(b)** MT-K748-ADP (colored) and microtubule unbound KIF14 motor domain structure (PDB: 4OZQ, semi-transparent gray) comparison. Structures are aligned to the KIF14 regions that interact with β-tubulin in the MT-KIF14-ADP structure. The KIF14 motor domain of the MT-K748-ADP structure is colored by the distance to equivalent Cα atoms in the MT-unbound structure (semi-transparent grey) according to the inset scale (in Å). **(c)** Displacement vectors (red arrows) between equivalent KIF14 motor domain Cα atoms in the MT-K748-ADP (light gray) and microtubule unbound (not shown) structures (alignment of structures as in b). **(d)** Nucleotide binding pocket comparison between microtubule unbound (semitransparent gray) and MT-K748-ADP (colored) structures. Both structures are aligned to their corresponding P-loops. **(e)** Superimposed MT-KIF14 complex structures (alignment as in b) in the ADP (pink) and apo states (blue). **(f)** Superimposed central β-sheet of all the MT-KIF14 complex structures (aligned to their corresponding P-loops). Numbers corresponds to the kinesin motor domain β-strands KS1 to KS8. Based on the amount of twist of the central β-strand the structures separate into two major groups, a more twisted group (blue and green) and a less twisted group (red). The low-twist structures (red) correspond to complexes in the closed conformation group and high-twist structures to the semi-closed (green), open and open* (blue) conformation groups. The orientation of the displayed structures relative to the microtubule is with the plus end up for (a), (b) and (d) and with the plus end away from the viewer (i.e. from the minus to the plus end) in (c) and (f).

Comparing the microtubule bound and unbound structures of KIF14 in the same ADP-bound nucleotide state also reveals a displacement between the tubulin interacting regions (Fig. 3b, c, Supplementary Movie 1). KIF14 helix-6 (KH6), which makes contacts with α-tubulin, moves relative to helix-4 (KH4), and KIF14 loops 8 and 12 (KL8, KL12) which make contacts with α- and β-tubulin (Fig. 3b-d). The displacement between these areas causes a movement between other regions of the KIF14 motor domain that to a first approximation can be described as a relative rotation between two sub-domains within the KIF14 head; a sub-domain that includes the regions that interact with β-tubulin and a sub-domain that includes KH6 which interacts with α-tubulin more towards the microtubule minus end and KIF14 helix-0 (KH0). Because the kinesin nucleotide binding pocket is located between these sub-domains their relative movement results in an altered nucleotide-binding-pocket architecture (Fig. 3d). From the microtubule unbound to the microtubule bound structure, the distance between the switch loops and the P-loop increases and there is a large displacement in the position of KH0 relative to the microtubule and the switch loops (Fig. 3d, Table 3). These structural rearrangements disrupt interactions between the ADP phosphate groups, the switch regions and the P-loop providing an explanation for the acceleration of product release associated with kinesin microtubule binding (Cross, 2016).

We found the structures of the motor domain in the ADP and apo states to be very similar in all the complexes investigated (Fig. 3e) and we refer to them as the open conformation group (Table 2, 3), according to their more open nucleotide binding pocket structure. The twist present in the central β-sheet of the KIF14 motor domain in the microtubule bound ADP and apo structures and in the microtubule unbound crystal structure is also very similar (Fig. 3f, Table 3) and higher than the one present in most kinesin crystal structures. This central β-sheet is another structural element common in related NTPases and based on comparison of myosin structures it is thought that the more twisted configuration represent an ADP-release/apo intermediate (Arora et al., 2014; Kull and Endow, 2013). The fact that the more twisted β-sheet is observed in the MT-KIF14-apo complex supports this interpretation.

There is a large displacement in the position of KH0 between the microtubule unbound and bound ADP and apo structures (Fig, 3b). We propose that the observed open KH0 position facilitates the incorporation of ATP into the nucleotide binding pocket or the exchange of ADP for ATP. This is consistent with molecular dynamics simulations that link movements of KH0, KL5 and SW1 with the incorporation of ATP into the nucleotide binding pocket (Hwang et al., 2017). Providing direct structural evidence for this proposal, we find that ATP analogues also readily bind to an open nucleotide binding conformation (open* structures, Tables 2 and 3, Figs. 5f, g, Supplementary Fig. 5 see, also section below: Ligands position in the open* motor domain conformation).

### ATP analogues induce closure of the KIF14 nucleotide binding pocket

The ATP analogue AMP-PNP and the ADP-Pi analogue ADP-AlF_x_ induce large conformational changes in the KIF14 constructs with at least 15 neck-linker residues (K748, K755 and K772), Table 2 Supplementary, Movie 1). Comparing the MT-K748-ANP and MT-K748-AAF with any of the apo or ADP structures reveals a large rotation of the KH0 containing sub-domain (Fig. 4a Supplementary, Movie 1). The MT-K748-ANP and MT-K748-AAF structures are very similar but not identical, indicating different responses of the KIF14 motor domain to whether the nucleotide in the binding pocket is in the pre- or post-hydrolysis state. In the ADP-AlF_x_ state, the motor sub-domain further away from the microtubule is slightly rotated relative to the position observed in the MT-K748-ANP complex (Fig. 4c Supplementary, Movie 1).

**Figure 4.**
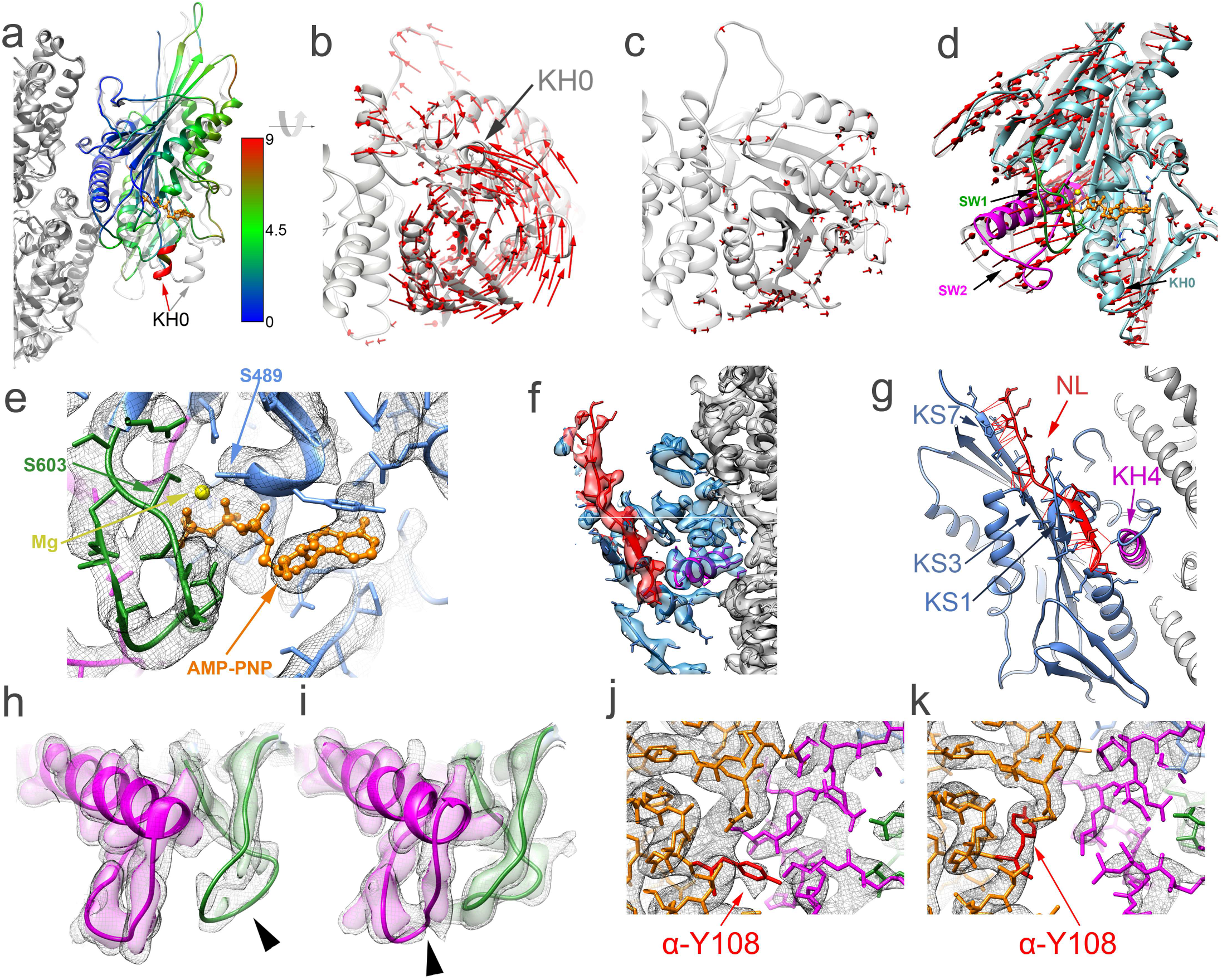
Closed nucleotide binding pocket conformation. **(a)** Superimposed MT-K748-ANP (colored) and MT-K748-ADP (semi-transparent gray) structures aligned to their corresponding tubulin β-chains. The KIF14 motor domain of the MT-K748-ANP structure is colored by the distance to equivalent Cα atoms in the MT-748-ADP structure according to the inset scale (in Å). **(b)** Displacement vectors (red arrows) between equivalent KIF14 motor domain Cα atoms in the MT-K748-ANP (light gray) and MT-K748-ADP (not shown) structures after alignment as in (a). **(c)** Displacement vectors (red arrows) between equivalent KIF14 motor domain Cα atoms in the MT-K748-AAF (light gray) and MT-K748-ANP (not shown). **(d)** Nucleotide binding pocket comparison with displacement vectors (red) between MT-K748-ADP (semitransparent gray) and MT-K748-ANP (colored) structures after aligning their P-loops. The KH4 and SW2 regions are colored magenta, the SW1 in green, and the AMP-PNP in orange. The rest of the motor domain is colored light blue. **(e)** Detail of the nucleotide binding pocket with corresponding densities (grey mesh) in the MT-K748-ANP structure. **(f)** Detail of the neck-linker domain with corresponding densities (semi-transparent colored surface) in the MT-K748-ANP structure. Neck-linker is colored red, the rest of the KIF14 motor in blue and tubulin in grey. **(g)** Neck-linker area in the MT-K748-ANP structure showing interacting residues with other areas of the motor domain. Pseudo-bonds between interacting atoms are represented as red lines (determined and displayed using the “find clashes and contacts” routine in UCSF-Chimera, see methods). **(h-i)** Switch loop regions in the MT-K748-ADP (i) and MT-K748-ANP (j) structures. SW1 in green and SW2 in magenta. Corresponding densities are showed at two iso-contour surfaces, higher (colored semi-transparent surface) and lower (grey mesh). Densities that disappear at the higher contour level (presumably more disordered) are pointed with the black arrowheads. **(j-k)** Detail of the microtubule-KIF14 interface of the MT-K748-ADP (j) and MT-K748-ANP (k) structures. Corresponding density iso-contour surface shown as a grey mesh. KIF14 SW1 and SW2 atomic models colored in green and magenta respectively and α-tubulin in orange except α-Y108 which is colored red. Note the distinct separation between α-tubulin and KIF14 residues and the alternate rotamer position of the α-Y108 side chain in the open ADP and closed AMP-PNP structures.

The rotation of the KH0 sub-domain going from the ADP to the AMP-PNP or ADP-AlF_x_ states is in the opposite direction to the one produced by microtubule binding in the apo or ADP states and larger in magnitude (Fig. 3b, c, Fig. 4a, b). It is also accompanied by a reduction in the twist of the central β-sheet (Fig. 3f, Table 3) and an opening of the hydrophobic pocket formed between KIF14 β-strand-1 (KS1) and helix-4 (KH4) that allows the neck-linker to dock onto the motor domain (Fig. 4f, g). The neck-linker is docked onto the motor domain establishing contacts with KS1 forming the cover-neck-bundle, a structural motif though to stabilize the neck-linker in the docked configuration and to be important for force generation (Hwang et al., 2008; Khalil et al., 2008). Further contacts at the tip of the motor domain are formed between the neck-linker and β-strands 3 and 7 (KS3 and KS7) (Fig. 4g).

In addition, the switch loops move towards the nucleotide in a direction opposite to the movement of the KH0 sub-domain to establish a more closed nucleotide binding pocket (Fig. 4d, e). There are also changes in the density associated with the tips of the switch loops that indicate changes in the mobility of these regions (Fig. 4h, i). In the ADP state the density at the tip of SW1 is relatively weak indicating higher mobility in this area compared to the tip of SW2. This situation is reversed in the AMP-PNP and ADP-AlF_x_ states.

The rotation of the KH0 containing sub-domain and the movement of the switch loops from the apo/ADP to the AMP-PNP/ADP-AlF_x_ conformations also induce structural rearrangements at the microtubule-KIF14 interface. KH6 changes orientation and the SW2 region moves away from α-tubulin (Fig. 4j, k, Supplementary Fig. 2). This rearrangement disrupts contacts between SW2 and α-tubulin Tyr-108 and this tubulin residue changes to an alternate rotamer conformation (Fig. 4j, k). Note that a salt bridge formed between conserved residues in SW1 and SW2 (Arg604 to Glu643 in KIF14), initially considered to be a defining characteristic of the closed catalytic configuration (Kull and Endow, 2002; Parke et al., 2010) also occurs in the open nucleotide configuration (Supplementary. Fig. 6). The switch loops, except the differences indicated above at their tips and near the microtubule interface, have similar topology and in all microtubule-bound structures.

The configuration of the motor domain around the nucleotide phosphate groups in the MT-K748 AMP-PNP and ADP-AlF_x_ states is similar to the closed ATP-hydrolysis-catalytic configuration, first observed in the microtubule unbound crystal structure of the kinesin-5 EG5 in the presence of AMP-PNP (Parke et al., 2010). This conformation of the KIF14 motor domain, together with the ones observed for the microtubule bound in the apo and ADP states and the microtubule unbound structure, define three major conformations of the KIF14 motor domain. Based on the corresponding nucleotide binding pocket configuration, we called these conformations closed, open, and semi-closed (Tables 2 and 3, Supplementary, Movie 1).

### Full neck linker docking is required for complete nucleotide binding pocket closure

Most models for the coordinated movement of kinesins along microtubules couple some step of the ATPase cycle with the position of the neck-linker domain (Andreasson et al., 2015; Hancock, 2016). However, how the position of the neck-linker may control the ATPase cycle is unclear. There is no direct structural data for any kinesin indicating how the position of the neck-linker could affect the structure of the nucleotide-binding-pocket. To investigate this issue, we obtained the cryo-EM structures of microtubules in complex with the constructs, K735 and K743 that have truncated neck-linkers (Fig.1b). We surmised that an absent or truncated neck-linker would mimic the situation when the neck-linker is mechanically pulled out from the docked configuration as it may occur under force or in a two-head microtubule bound intermediate.

In the apo and ADP states the motor domain structures are very similar for all the complexes investigated; this is regardless of whether the KIF14 construct contained a truncated or a full neck-linker, they all conform to the open configuration (Fig. 3e, Tables 2, 3). On the other hand, in the presence of AMP-PNP or ADP-AlF_x_ the motor domain structures of the MT-K735 and MT-K743 complexes do not resemble the closed configuration observed with constructs that include a full neck-linker (Tables 2, 3).

The motor domain in the MT-K735-ANP complex shows a new configuration where the KH0-containing-sub-domain is rotated to an intermediate position between the open and closed configurations (Fig. 5a, Supplementary Fig. 3), but as in the open configuration the switch loops remain further away from the P-loop (Fig. 5b). In the MT-K735-ADP-AlF_x_ state, the whole motor domain structure closely resembles the open conformation observed in the apo/ADP states (Fig. 5a, b). These results show that without the neck linker (or with just two residues of it), the following observations can be made: 1) ATP analogues with the γ-phosphate group covalently linked such as AMP-PNP induce a partial rotation of the KH0-sub-domain towards the closed configuration; 2) the post-hydrolysis analogue ADP-AlF_x_ does not induce any sub-domain rotation and the motor domain mostly resembles the open configuration; 3) neither analogue induces the closed configuration of the motor domain observed with the construct that include at least 15 neck linker residues.

**Figure 5.**
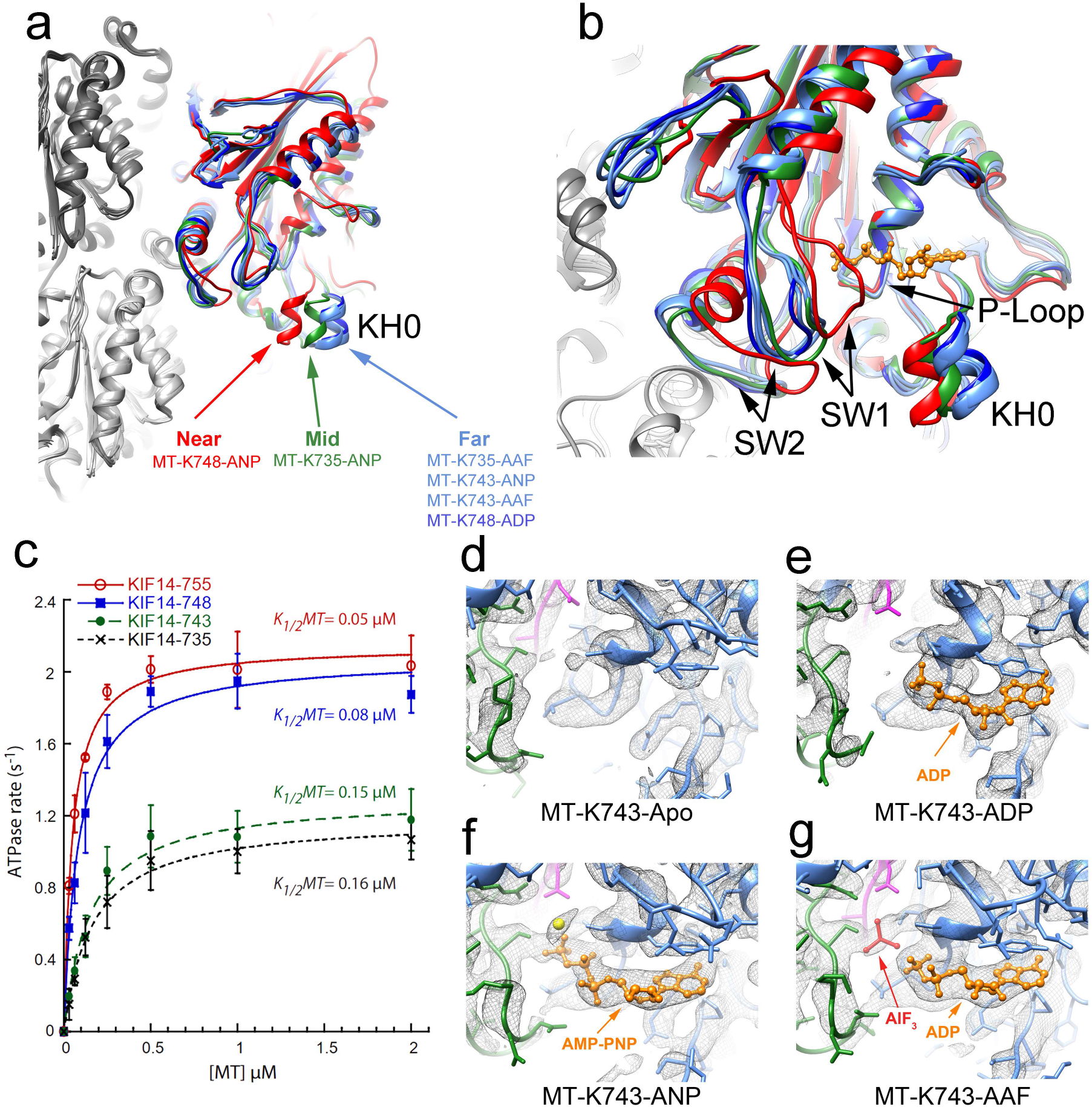
Neck linker control of nucleotide binding pocket closure. **(a)** Superimposed MT-K735-ANP, MT-K735-AAF, MT-K743-ANP, MT-K743-AAF, MT-K748-ANP, MT-K748-ADP and MT-K748-ANP structures. All the structures are aligned to their corresponding β-tubulin chains. The near position of the KH0 sub-domain observed in the MT-K748-ANP closed structure (red) is not observed in the constructs with truncated neck-linkers K735 and K743. In the MT-K35-ANP structure (green) the KH0 subdomain adopts an intermediate position. In the MT-K735-AAF, MT-K743-ANP, MT-K743-AAF structures (blue) the KH0 sub-domain adopts a far position similar to the MT-K748-ADP open structure (blue). **(b)** Nucleotide-binding-pocket view showing the same structures with the same colors as in (a) superimposed and aligned to their corresponding P-loops. The switch loops, SW1 and SW2 and the KH0 subdomain are positioned farther away from the P-loop and the nucleotide in the MT-K735-ANP, MT-K735-AAF, MT-K743-ANP and MT-K743-AAF than in the MT-K748-ANP closed structure (red). The positions of SW1, SW2 and KH0 relative to the P-loop of the K735 and K743 constructs instead resemble the ones in the MT-K748-ADP open structure (blue). **(c)** Microtubule stimulated ATPases of K735, K43, K748 and K755 constructs. **(d-g)** Nucleotide-binding-pocket detail of MT-K743-apo, MT-K743-ADP, MT-K43-ANP and MT-K743-AAF cryo-EM maps and fitted models. Cryo-EM densities shown as an iso-surface grey mesh, KIF14 model in blue with SW1 in green, SW2 in magenta, bound nucleotide (ADP or AMP-PNP), AlF_3_ and Mg^2+^ in orange, red and yellow respectively.

The structures of the MT-K743 complex in the AMP-PNP and ADP-AlF_x_ also do not conform to the closed configuration observed with the constructs that include a full neck-linker. In both cases the motor domain structure is best fitted to an open configuration model and a docked neck-linker is not resolved (Figs 2, 5a, b). However, different from the MT-K735 complexes, the corresponding cryo-EM density maps also show weaker densities spreading towards the closed conformation position suggesting the presence of a small fraction of the motor domains in the closed conformation (< 17 %) of the population (Supplementary Fig. 4). The lack of a docked neck linker conformation in the K743 construct is surprising considering that it contains the residues that form the cover-neck-bundle and the asparagine latch (N743 in KIF14), thought to hold the neck-linker in the docked conformation (Budaitis et al., 2019; Hwang et al., 2008). The fact that the K743 construct cannot adopt the docked configuration but the K748 can, shows that the additional contacts formed by the extra five residues of the 748 construct (Fig. 4g) are required to stabilize the docked configuration and that a near full neck-linker from the end of KH6 to the beginning of the CC1 coiled coil domain needs to be docked for the nucleotide-binding-pocket to be fully closed.

Taking together the results with the K735 and K743 constructs show that without a fully docked neck-linker the KIF14 motor domain cannot adopt the closed catalytic conformation and instead adopt a conformation most similar but not identical to the open structures observed in the apo and ADP states. These structures can be grouped into a fourth group, open* (Tables 2 and 3). Within this group the most distinctive conformation of the motor domain corresponds to the intermediate configuration observed in the MT-K735-ANP complex (Fig. 5a, b, Tables 2 and 3).

Consistent with the structural results, we found that the K735 and K743 constructs have reduced microtubule stimulated ATPase activity relative to the longer KIF14 constructs (Fig. 5c). Thus, the neck-linker position and the ATPase activity of the motor domain are fully coupled. ATP binding induces closure of the nucleotide binding pocket and neck-linker docking (as shown in the MT-K748-ANP structure). Conversely, preventing neck-linker docking impedes full closure of the nucleotide binding pocket and inhibits the ATPase activity.

### Unique ligands position in the open* motor domain conformation

The MT-K735 and MT-K743 maps in the AMP-PNP and ADP-AlF_x_ states show clear nucleotide associated densities in the active site indicating that both nucleotide analogues bind to the open* configuration of the motor domain (Fig. 5d-g, Supplementary Fig. 5 Supplementary Movie 1). In the case of the ADP-AlF_x_ maps, the ligands adopt a distinct configuration different from the one observed in the closed nucleotide binding pocket structures. Both the MT-K735-AAF and the MT-K743-AAF maps show two separated densities in the nucleotide binding pocket that can be attributed to the ADP and AlF_x_ groups (Fig. 5g). These densities are not present in the apo map (Fig. 5d) and the extra density attributed to the AlF_x_ group is absent in the ADP map (Fig. 5e). Also, there is no density that could be attributed to the Mg^2+^ ion as it is the case in the AMP-PNP maps (Figs. 4e and 5f). Fitting the ADP and AlF_x_ ligands into the densities places the AlF_x_ and ADP groups in the MT-K735-AAF and the MT-K743-AAF complexes ~2 Å further separated than in the MT-K748-AAF complex. To our knowledge, this is the first indication of an alternative configuration of the ligands mimicking the hydrolysis products in the kinesin active site. The lack of the Mg^2+^ coordinating atom and the increased separation between the nucleotide and the γ-phosphate group mimic indicates a weakened interaction of this group in the active site that may facilitate its release. This suggests that that neck-linker undocking (e.g. under tension) and the consequent opening of the nucleotide binding pocket may promote Pi release from the active site after ATP hydrolysis.

### One- and two-head microtubule bound KIF14 structures

The structure of the core motor domain of the MT-K755 and MT-K772 complexes in the apo and ADP states conform to the open configuration, as it is also the case for the complexes with the shorter KIF14 constructs (Fig. 3e, Tables 2 and 3). However, different from the MT-K743-ADP/apo and MT-K748-ADP/apo maps the MT-K755-ADP/apo and the MT-K772-ADP/apo show a neck-linker associated density going sideways reaching the motor domain of an adjacent protofilament (Figs. 2, 6a, b). Extra densities that could be associated with a partner motor domain in the dimeric K772 construct are not observed, suggesting that the partner head is not in a well-defined location. Another possibility to explain the lack of a distinct density associated with the partner motor domain would be that the two motor domains are bound to adjacent protofilaments in equivalent configurations. However, if this were the case two alternative neck-linkers orientations would be expected, one as observed and another going from helix-6 in the motor domain (where the neck-linker starts) to the dimerization domain at the tip of the observed neck-linker. Because such density is not observed, the map is better fitted with a model in which the KIF14 dimer in the apo and ADP states binds to the microtubule in a one-head bound state. In this state, one head is bound to the microtubule with the nucleotide binding pocket in the open configuration and the other is tethered and presumably in a semi-closed configuration similar to the microtubule unbound KIF14 crystal structure (K-ADP_4OZQ, Table 2).

**Figure 6.**
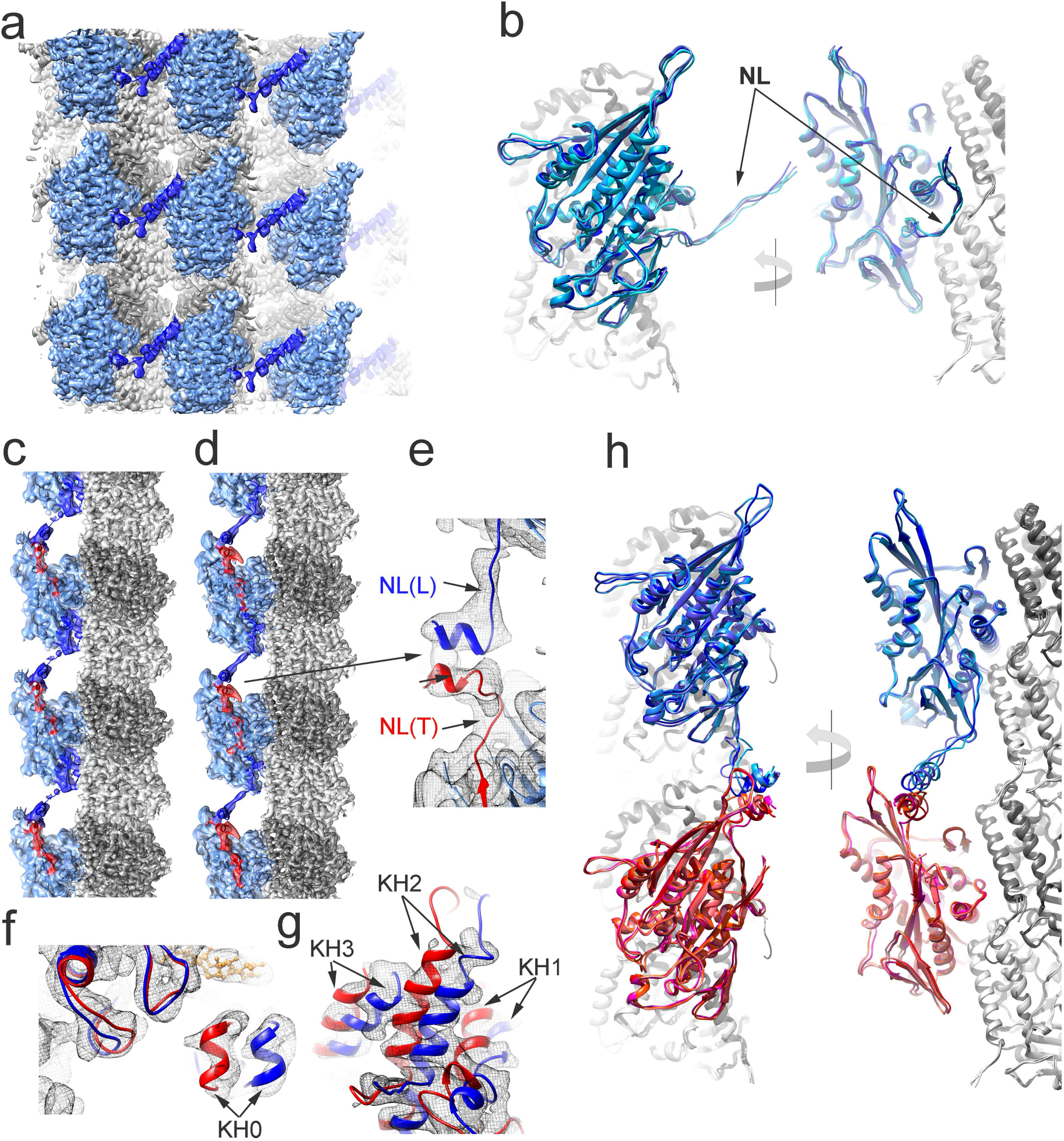
Microtubule bound KIF14 dimer structures. **(a)** Iso-surface representation of nine adjacent asymmetric units of the MT-KIF14-ADP cryo-EM map colored according to the fitted atomic model regions. The KIF14 motor domain and neck-linker are colored light and dark blue respectively. α- and β-tubulin regions colored light and dark grey respectively in panels a, b, c, d and h. **(b)** Superimposed MT-K755-ADP, MT-K755-apo, MT-K772-ADP and MT-K772-apo structures. All the structures aligned to their corresponding β-tubulin chains. The KIF14 motor domain of each model is colored in a different shade of blue. NL: Neck-linker. **(c)** MT-K772-ANP cryo-EM map. **(d)** MT-K755-ANP cryo-EM map. c-d show three asymmetric units of an isolated protofilament. The KIF14 motor domain is colored blue with docked and undocked neck-linker regions colored red and dark blue respectively. **(e)** Detail of the connecting neck-linkers and part of the coiled coil region of trailing (red) and leading KIF14 motor domain structures in the MT-K755-ANP complex inside the corresponding density (grey mesh). **(f-g)** MT-K755-ANP cryo-EM (grey mesh) with fitted open (blue) and closed (red) conformations of the KIF14 motor domain. (f) shows the region near kinesin motor domain helix 0 (KH0) and (g) the back of the motor domain further away from the microtubule with kinesin motor domain helices 1 2 and 3 (KH1, KH2, KH3). **(h)** Superimposed KIF14 dimeric structures in the MT-K755-ANP, MT-K755-AAF, MT-K772-ANP and MT-K772-AAF structures. Leading and trailing KIF14 motor domains colored blue and red respectively. Structures oriented in all the panels with the microtubule plus end up.

Different from the apo and ADP states in the AMP-PNP and ADP-AlF_x_ states the MT-K755 and MT-K772 complexes show a density connecting the motor domains along a protofilament (Figs. 2, 6c-e). This density can be well fitted to the undocked neck-linker of a motor domain positioned further towards the microtubule plus end (leading head) and part of the coiled coil domain connecting with the docked neck-linker of a motor domain positioned more towards the microtubule minus end (trailing head) (Fig. 6e). Thus, the connecting density between motor domains indicate the presence of a leading and trailing two-head-bound intermediate.

In the helically averaged cryo-EM maps the densities corresponding to the leading and trailing heads would be averaged, and in fact the maps indicate the presence of two distinct structures averaged. The densities in the motor area can be very well fitted with coexisting open and a closed motor domain structures (Fig. 6f, g, Supplementary Fig. 4). Thus, the cryo-EM data shows that the KIF14 dimer in the AMPP-PNP and ADP-AlF_x_ states binds to the microtubule in a two-head bound configuration with leading and trailing heads in distinct conformations (Fig. 6h). We assigned the open and closed conformations to the leading and trailing head respectively. This is based on the results with the shorter constructs indicating that an undocked or truncated neck-linker always results in an open conformation, regardless of the nucleotide present in the active site, and that the closed conformation is only achieved with a 15 residues fully docked neck linker, as shown by the MT-K748-ANP/AAF structures (Table 3).

There is some uncertainty regarding the precise structure of the undocked neck-linker and coiled coil domain because the density and achievable resolution of this part of the map is lower, particularly for the K772 construct. This is likely the result of increased structural flexibility and mobility in this area. However, we are confident that the densities in this area are produced by an undocked neck-linker and coiled coil structure as modeled because the densities are well accounted by the model and they appear only in the longer K755 and K772 constructs (Fig. 2). A density connecting the two-motor connection is not observed in the maps with construct K748 which is truncated just before the start of the CC1 domain.

Similar dimeric like structures were observed with the K755 and K772 constructs, even though only the K772 constructs is dimeric in solution (Fig. 1c). Construct K755 does not include the full CC1 domain but includes part of the first heptad repeat. Our results show that this incomplete heptad, although insufficient to induce dimerization in solution (Fig. 1c), is able to do so when many motor domains are bound in close proximity in the microtubule lattice.

The cryo-EM-maps also provide evidence that the nucleotide-binding-pocket of both the leading and trailing head is occupied by the ATP like analogues. This is contrary to the expectation that ATP binding to the leading head in a two head bound intermediate would be prevented (Dogan et al., 2015; Rosenfeld et al., 2003; Uemura and Ishiwata, 2003). If nucleotide binding to the leading head would be prevented it would be expected the nucleotide associated density to be half of what it is observed in the constructs that do not form dimers. Instead, the nucleotide associated densities of the MT-K772 and MT-K755 constructs in the AMP-PNP and ADP-AlF_x_ states are similar or higher than in the cryo-EM maps of the complexes with the constructs lacking the dimerization domain (K735, K743, K748) and that do not contain leading and trailing heads averaged (Supplementary Fig. 5). Also, the fact that a density associated with the added nucleotides is found in all the cryo-EM maps, regardless if the motor domain is in the closed or the open configuration, further supports the conclusion that both heads of the two head bound intermediate have a bound nucleotide in the active site.

## Discussion

We have solved the near atomic resolution cryo-EM structures of five distinct KIF14 protein constructs bound to microtubules, in four nucleotide conditions mimicking key steps of its ATPase cycle. Our data provides a comprehensive view at the highest resolution available of the mechano-chemical cycle of a motile kinesin and provide the structural basis for a mechano-chemical model for kinesin plus-end translocation (Fig. 7, Supplementary Movie 2). In the following paragraphs, we further discuss the conclusions that can be drawn from the data and how they relate to previous studies and models of kinesin motility.

**Figure 7.**
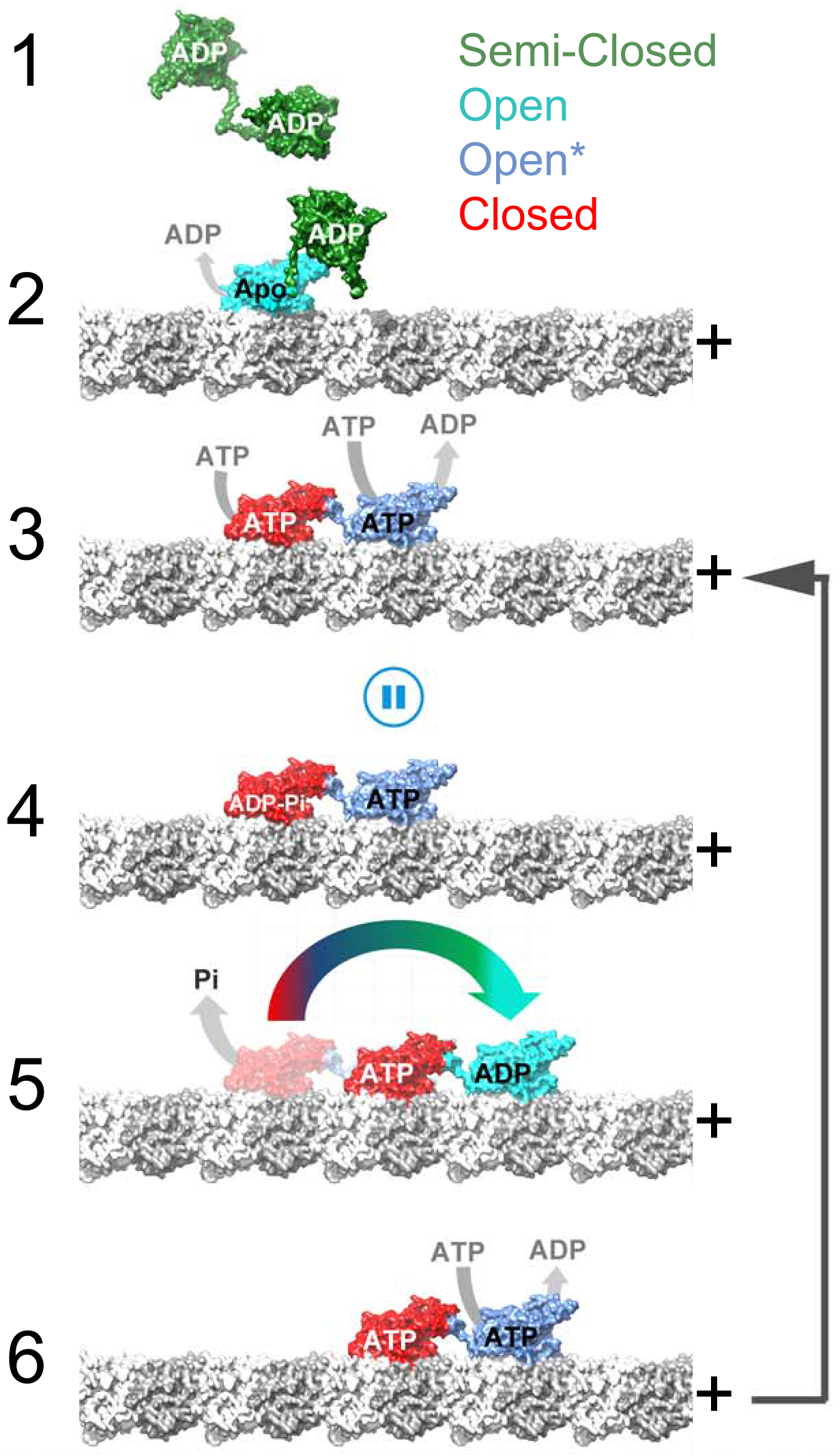
Kinesin dimer coordinated mechano-chemical cycle model. The four major KIF14 motor domain conformations determined, semi-closed, open, open* and closed (Table 3) are colored green, cyan blue and red respectively. α- and β-tubulin are colored light and dark grey respectively and the microtubule protofilaments are oriented with the plus end to the right. Starting from a microtubule unbound semi-closed-ADP conformation (state 1) binding to the microtubule induces opening of the nucleotide-binding pocket and release of ADP (state-2). In this state the neck-linker of both motor domains is undocked preventing the tethered motor domain from reaching the next tubulin binding site. The tethered motor domain remains mobile (disordered) and the dimer adopts a one-head bound conformation similar to the MT-K755 – apo/ADP and MT-K772-apo/ADP structures. Opening of the nucleotide pocket in the microtubule-bound head induces ADP release and allow ATP binding. ATP binding induces closure of the nucleotide binding pocket and neck-linker docking. The docked neck-linker of the bound head positions the accompanying, tethered head, in a leading position (towards the microtubule plus end). The now leading head binds to the microtubule with its neck-linker undocked and pulled backward by the microtubule bound trailing head. A two-head microtubule bound intermediate similar to the MT-K755-ANP and MT-K772-ANP structures is formed (state 3). This two head intermediate establish a coordinating gate in the chemo-mechanical cycle of the two motor domains. The undocked neck-linker in the leading head prevents it from adopting the closed catalytic conformation pausing hydrolysis until the trailing head releases from the microtubule. ATP hydrolysis is then favored in the neck-linker docked trailing head but not in the leading head leading to state 4. We speculate that in this state inter-head tension may also cause partial undocking of the trailing head neck-linker leading to an open* trailing head configuration (similar to the MT-K735-AAF and MT-K743-AAF structures) where product release is facilitated. Hydrolysis and/or product release in the trailing head leads to detachment from the microtubule and a transient one-head bound intermediate is formed (state 5). In the one head bound intermediate the neck-linker of the ATP bound leading head docks and places the tethered head forward toward the plus end. The two motor heads change leading and trailing positions and the now microtubule bound leading head in the open* configuration releases ADP and binds ATP (state 6). State 6 is similar to 3 but with the two heads exchanging leading and trailing positions and the whole molecule displaced 8 nm in the plus direction.

### Conformational changes induced by microtubule binding

The microtubule bound KIF14 open structures share some similarities with the near atomic resolution crystal structure of kinesin-1 bound to curved tubulin in the apo state (Cao et al., 2014). In both cases, the nucleotide binding pocket becomes more open relative to the tubulin unbound structure. but there are also significant structural differences between the complexes. In the kinesin-1-tubulin apo structure, the switch-1 loop is mostly disordered and becomes ordered in the presence of ADP-AlF_x_ (Cao et al., 2014; Gigant et al., 2013). On the other hand, in all the KIF14-microtubule structures, including the apo state, this loop is mostly ordered and forms the salt bridge with switch-2 (Supplementary Fig. 6) that was previously thought to be unique to the catalytically active closed conformation. The relative position between tubulin interacting regions and other areas of the motor domain are also slightly different (Supplementary Fig. 7). Similar differences are also noted when comparing the KIF14-microtubule complex structures and models derived from sub-nanometer resolution cryo-EM structures of the kinesin-1 microtubule complex (Shang et al., 2014). How much of these differences are due to the different kinesin types (Kinesin-3 KIF14 vs. Kineins-1 KIF5B), different tubulin structures (curved vs. straight) or the resolution of the cryo-EM structures is not fully clear yet. To address this issue, structural studies of other kinesin-microtubule complexes at similar or higher resolution than achieved here are needed. Although there are several cryo-EM structures of kinesin microtubule complexes available, only recently it has become possible to achieve the resolution needed (< 4 Å in the kinesin part of the map) to fully trace the polypeptide chain and to resolve the positions of side chains, the nucleotide species in the active site and the conformation of exposed and potentially mobile regions such as the switch loops and the neck-linker.

### The KIF14 neck-linker controls nucleotide-induced conformational changes

There is good evidence indicating that binding of ATP or its analogues to the motor domain of microtubule-bound plus-end-directed-motile-kinesins, induce neck-linker docking (Asenjo et al., 2006; Rice et al., 1999; Tomishige and Vale, 2000) and closure of the nucleotide binding pocket (Atherton et al., 2014; Kikkawa and Hirokawa, 2006; Shang et al., 2014). However, whether and how the neck-linker position could allosterically control the conformation of the motor domain and its catalytic activity was not clear. Our results show that without a docked neck-linker, the motor domain is prevented from adopting the closed catalytic conformation (Fig. 5, Tables 2 and 3) and provide the structural basis for a fully reciprocal relationship between the neck-linker position and the nucleotide-binding-pocket. Closure of the nucleotide binding pocket induces neck-linker docking and conversely, preventing neck-linker docking impedes the nucleotide binding pocket from adopting the fully closed catalytically active conformation. Our results also suggest that neck linker undocking after ATP hydrolysis promote Pi release from the nucleotide binding site.

The fact neck-linker docking was not observed for constructs shorter than the K748 construct also shows that near the full length of the neck-linker (15 residues from R734 to A748) needs to be docked for the nucleotide binding pocket to fully close. This and the reciprocal relationship between neck-linker docking and nucleotide binding pocket structures appear to be perfectly optimized for coordinated plus end directed movement. A fully docked neck-linker would place the partner motor domain of the KIF14 dimer in the leading position, close to its next binding site along the protofilament. At the same time the 8 nm distance between binding sites could only be covered with the neck-linker of the leading head in an undocked position. Thus, in a two-head bound state a docked neck-linker in the trailing motor domain and undocked neck-linker in the leading head would keep their catalytic cycles out of step as only the nucleotide-binding-pocket of the trailing head would be in the closed catalytic configuration.

It is plausible that the number of residues required to stabilize the docked conformation varies between kinesins as the neck-linker sequence is not highly conserved. This may provide a mechanism for fine tuning the catalytic activity of the motor, its response to tension or the coordination between heads in a dimer. As shown here for KIF14, ten or less residues of the neck linker (as in the K743 and shorter constructs) are insufficient to fully stabilize the docked conformation. In contrast, for the non-motile kinesin-13 KLP10A only six neck-linker residues are sufficient to stabilize a docked conformation and a closed nucleotide binding pocket when the motor domain is bound to curved tubulin (Benoit et al., 2018).

The KIF14-microtubule complex structures also show that the step of the ATPase cycle that is regulated by the position of the neck-linker is hydrolysis rather than nucleotide binding, as has been proposed based on single molecule force mechanics and kinetic experiments (Dogan et al., 2015; Rosenfeld et al., 2003; Uemura and Ishiwata, 2003). If nucleotide binding were prevented when the neck-linker is forced out of the docked configuration, then an empty nucleotide binding pocket would be expected in the KIF14 constructs with truncated neck-linkers (K735 and K743). Instead, the nucleotide present in the solution, ADP, AMP-PNP or ADP-AlF_x_, is always found in the nucleotide binding pocket of these constructs (Fig. 5e-g, Supplementary Fig. 5). Furthermore, the data also indicate that nucleotide binds to both the leading and trailing heads of the two head microtubule bound intermediates formed by the K755 and K772 constructs in the AMP-PNP and ADP-AlF_x_ states (Supplementary Fig. 5). The capacity of the kinesin motor domain to bind nucleotides regardless of the position of the neck-linker is unlikely to be a feature unique to KIF14. A previous cryo-EM study of microtubule bound kinesin-1 dimers also suggested that nucleotides were present in both the leading and the trailing heads (Liu et al., 2017). In addition, a recent structure of another kinesin (kinesin-13), where the nucleotide binding pocket is prevented from closing by a different mechanism, also show an ATP analogue bound in the active site (Benoit et al., 2018). Thus, controlling closure of the nucleotide binding pocket and entering the hydrolysis competent step appears to be a general regulatory mechanism in the kinesin super-family. Although we cannot rule out small changes in nucleotide affinity, we do not expect them to be a limiting factor at physiological molar concentrations of ATP, similar to the ones used in this study.

### Leading and trailing structures of the KIF14 dimer

The longer KIF14 constructs that includes part of the coiled coil dimerization domain, K755 and K772, showed for the first time at near atomic resolution what the structures of a kinesin dimer two-head microtubule bound intermediate looks like. Such intermediate is present in most models of kinesin plus-end directed processive movement and it is likely the dominant state during translocation at physiological ATP concentrations (Asenjo et al., 2003; Asenjo and Sosa, 2009; Guydosh and Block, 2009; Hancock, 2016). The structures of the MT-K755 and MT-K772 complexes with AMP-PNP or ADP-AlF_x_ shows two microtubule bound coexisting conformations of the kinesin motor domain providing evidence of coordination between the two motor domains. The ATPase cycle of the two motor domains is kept out of step as the undocked neck-linker of the leading head keeps the catalytic site in the open configuration inhibiting ATP hydrolysis (Fig. 7 steps 3 to 4). This control mechanism where the ATPase cycle of the leading head is slowed down by the backward pulling rear head has been referred as front-head-gating (Andreasson et al., 2015) and our data provides direct structural evidence for it. However, distinct from previously proposed models, our data argue that the step of the ATPase cycle that is inhibited in the leading head is ATP hydrolysis rather than nucleotide binding (see previous section).

The dimeric K772 constructs in the apo and ADP states also provide a high-resolution view of the one-head microtubule bound intermediate occurring during kinesins translocation when one kinesin motor domain changes binding sites along the microtubule. In this intermediate, the bound motor domain is in the open configuration with an undocked neck-linker pointing to the side and the partner motor head is mobile and not visible in the cryo-EM maps (Fig. 4b). This fits well with data indicating that kinesin-1 dimers during translocation form a transient one-head microtubule bound intermediate with a highly mobile tethered head in the so called ATP-waiting state (Asenjo and Sosa, 2009; Guydosh and Block, 2009; Mori et al., 2007). The conformation of the neck-linker in the K772 and K755 apo/ADP structures indicate that the tethered head is positioned sideways and toward the microtubule plus end (the direction of movement) relative to the trailing position seen in the two-head microtubule bound intermediate.

In summary, we obtained the near-atomic resolution structures of eighteen distinct kinesin KIF14 microtubule complexes providing the structural basis for a coordinated translocation mechanism where the hydrolysis step of each motor domain is controlled by the conformations of their connecting neck-linkers domains (Fig. 7). Given the high degree of motor domain homology and that KIF14 shares a similar quaternary structure with other plus-end directed motile kinesins, most aspects of the proposed model are likely to be generally applicable to this group of motors.

## Acknowledgments

This work was supported by NIH grant R01GM113164 (HS), Canadian Cancer Society Research Institute Grant #703405 (BHK) and Natural Sciences and Engineering Research Council of Canada Discovery Grant (BHK). B.H.K. is a recipient of the Fonds de Recherche du Québec– Santé (FRSQ) Chercheure-Boursière Junior 1, Junior 2 Awards and the Canadian Institutes of Health Research (CIHR) New Investigator Award. Cryo-EM data collection was performed at the Simons Electron Microscopy Center and National Resource for Automated Molecular Microscopy located at the New York Structural Biology Center, supported by grants from the Simons Foundation (SF349247), NYSTAR, and the NIH National Institute of General Medical Sciences (GM103310) with additional support from Agouron Institute (F00316) and NIH (OD019994). We thank Laura Yen, Misha Kopylov, Daija Bobe, Ed Eng and Bill Rice for assistance and support during data collection. We thank The Albert Einstein College of Medicine (AECOM) Analytical Imaging Facility for electron microscopy support and the AECOM High Performance Computing Facility for computing support.

---

**Supplementary Table 1.** Additional cryo-EM data collection and refinement parameters.

| **Dataset** | **Microscope used** | **Pixel Size (Å)** | **Box size (pixels)** | **Magnification anisotropy % <sup>a</sup>** | **Magnification anisotropy angle <sup>a</sup>** | **Per particle CTF refinement** | **15R particle cleaning** |
| --- | --- | --- | --- | --- | --- | --- | --- |
| 735-ANP | Krios 1 | 1.073 | 512 | n.c. | n.c. | N | N |
| 735-AAF | Krios 1 | 0.831 | 664 | 0.33 | 88.7 | Y | N |
| 735 apo | Krios 1 | 1.073 | 512 | n.c. | n.c. | N | N |
| 743-ANP | Krios 2 | 0.849 | 664 | 1.45 | 1 | Y | Y |
| 743-AAF | Krios 3 | 0.825 | 664 | 0.54 | 48.4 | Y | Y |
| 743-ADP | Krios 1 | 0.828 | 664 | 1.06 | 87.7 | Y | Y |
| 743 apo | Krios 2 | 0.848 | 664 | 1.69 | 179 | Y | Y |
| 748-ANP | Krios 1 | 0.828 | 664 | 1.07 | 89.8 | Y | Y |
| 748-AAF | Krios 3 | 0.825 | 664 | 0.52 | 47.4 | Y | Y |
| 748-ADP | Krios 2 | 0.849 | 664 | 1.46 | 175.9 | Y | Y |
| 755-ANP | Krios 1 | 1.072 | 512 | n.c. | n.c. | Y | Y |
| 755-AAF | Krios 2 | 0.848 | 664 | 1.6 | 179.2 | N | N |
| 755-ADP | Krios 1 | 0.831 | 664 | 0.35 | 70 | N | N |
| 755 apo | Krios 2 | 1.088 | 512 | 1.5 | 171.7 | N | N |
| 772-ANP | Krios 1 | 1.073 | 512 | n.c. | n.c. | N | N |
| 772-AAF | Krios 3 | 0.826 | 664 | 0.44 | 49.6 | Y | Y |
| 772-ADP | Krios 2 | 0.849 | 664 | 1.46 | 3.6 | N | N |
| 772 apo | Krios 2 | 0.850 | 664 | 1.25 | 3.1 | N | N |

**Notes**
<sup>a</sup> Given when estimated and corrected for a given dataset. n.c.: magnification anisotropy not corrected.

**Supplementary Fig. 1.**
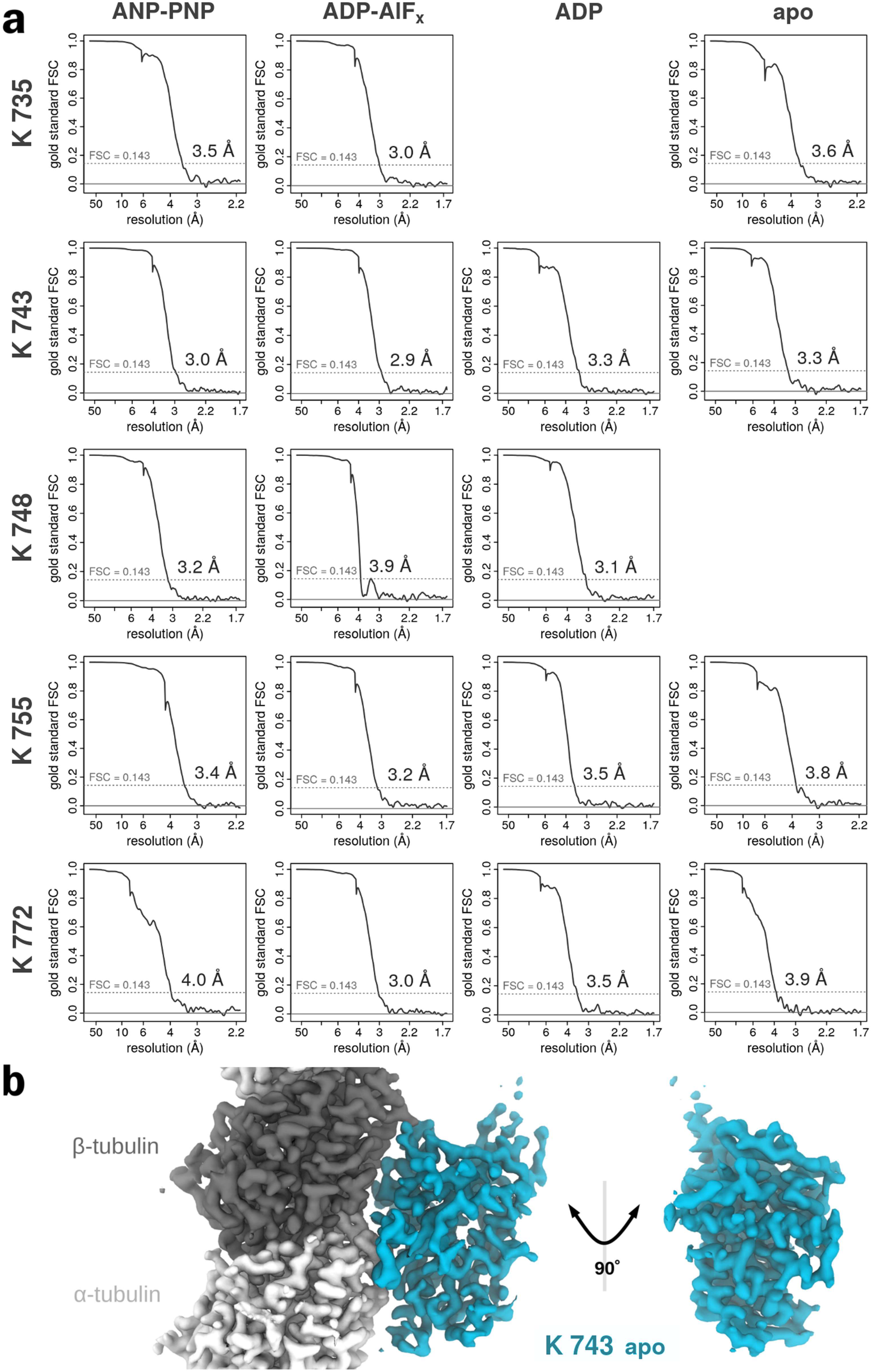
Fourier Shell Correlation Curves (FSC). **(a)** FSC curves for each of the eighteen structures solved with the estimated resolution (FSC_0.143_) indicated. **(b)** Iso-density surface representation of the MT-K743-apo cryo-EM map. Note that the polypeptide chain, the helical paths of the α-helices and most side chains are well resolved, as expected from the estimated resolution of the maps.

**Supplementary Fig. 2.**
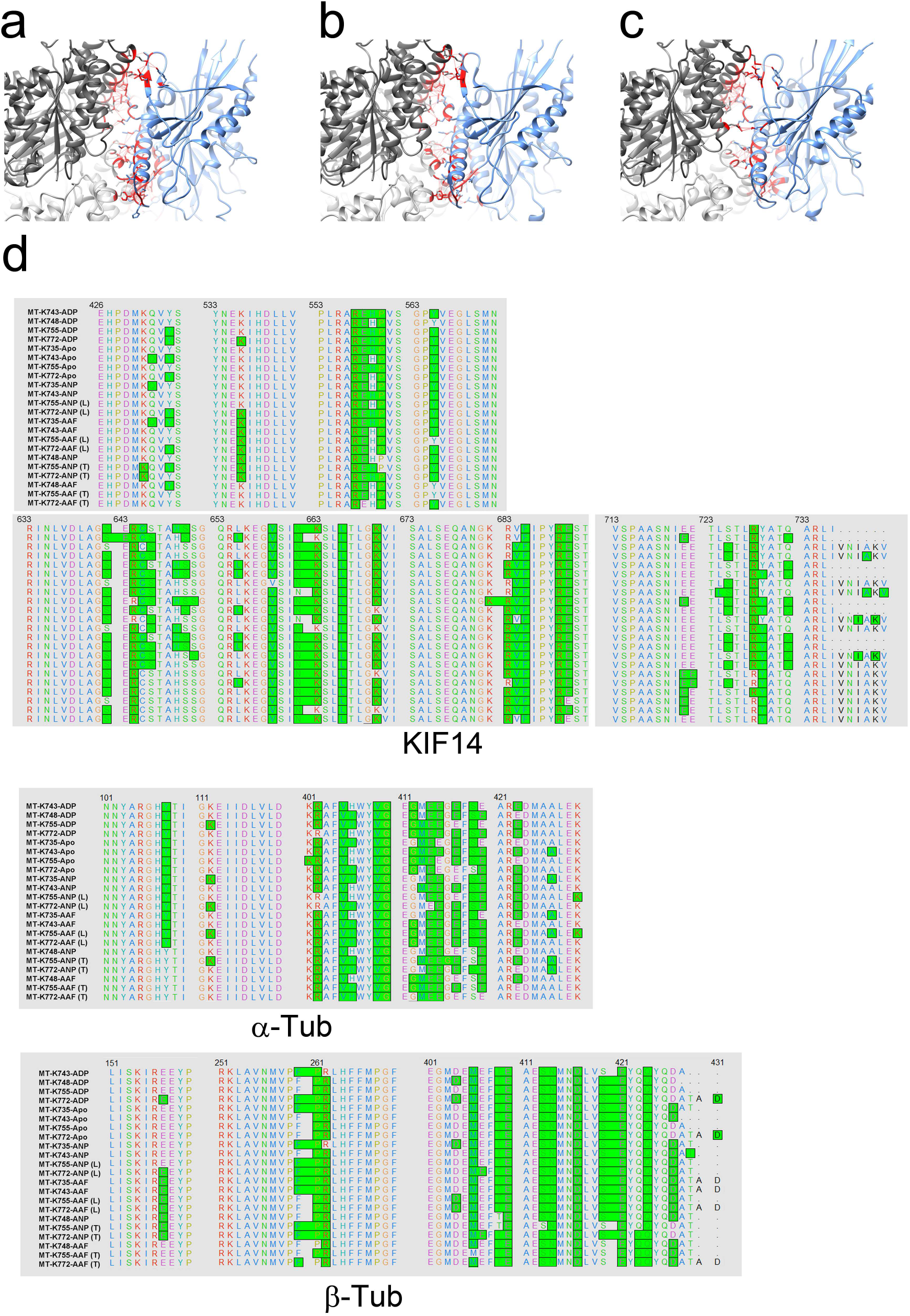
KIF14-Microtubule interacting residues. **(a-c)** Interface between the KIF14 motor domain and the microtubule highlighting interacting atoms as red lines (pseudo-bonds). Interacting atoms at the interface were determined and displayed using the find-clashes- and-contacts routine in UCSF-Chimera. Three selected interfaces are shown corresponding to the MT-K743-apo (a), MT-K748-ADP (b) and MT-K748-ANP structures (c). **(d)** KIF14 α- and β-tubulin residues identified as interacting residues in each microtubule-KIF14 complex are highlighted in green in the displayed amino-acid sequences.

**Supplementary Fig. 3.**
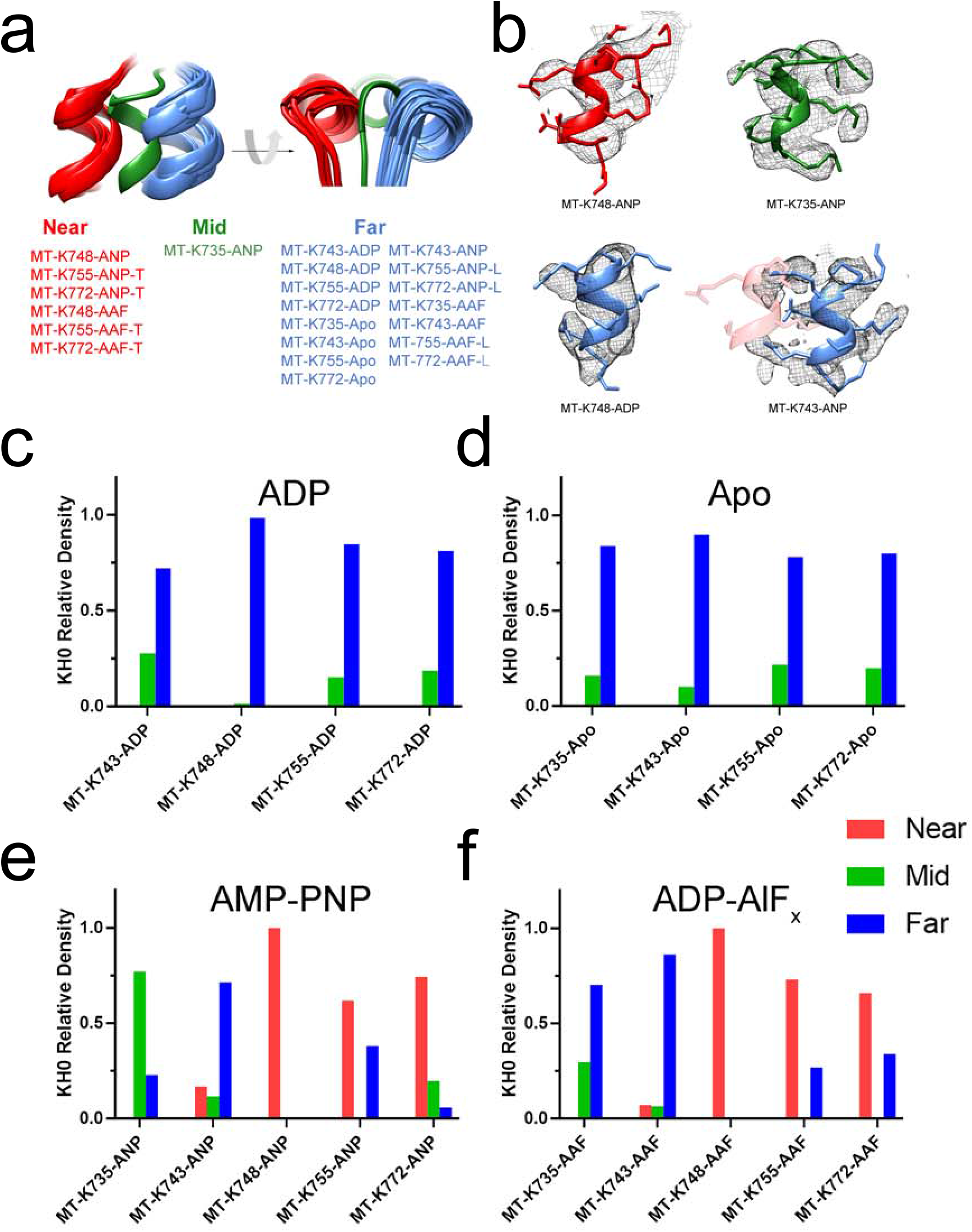
Cryo-EM map densities at distinct KIF14 helix 0 (KH0) locations. **(a)** Three distinct locations of KH0, near (to the microtubule), mid and far are identified after aligning all the models to their corresponding β-tubulin subunit. **(b)** KH0 associated densities in four selected cryo-EM maps. Note that the KH0 associated densities are well localized in the top and left structures while in the bottom right (MT-K743-ANP) they spread from the far (blue) to the closed position (semi-transparent red). **(c-f)** Relative densities associated with each of the three KH0 locations for all the cryo-EM maps. Relative densities were calculated as RD_x_ = D_x_ / (D_N_ + D_M +_ D_F_), where Dx corresponds to the average density minus background at each of the three locations, Near (N), Mid (M) and Far (F) (see methods).

**Supplementary Fig. 4.**
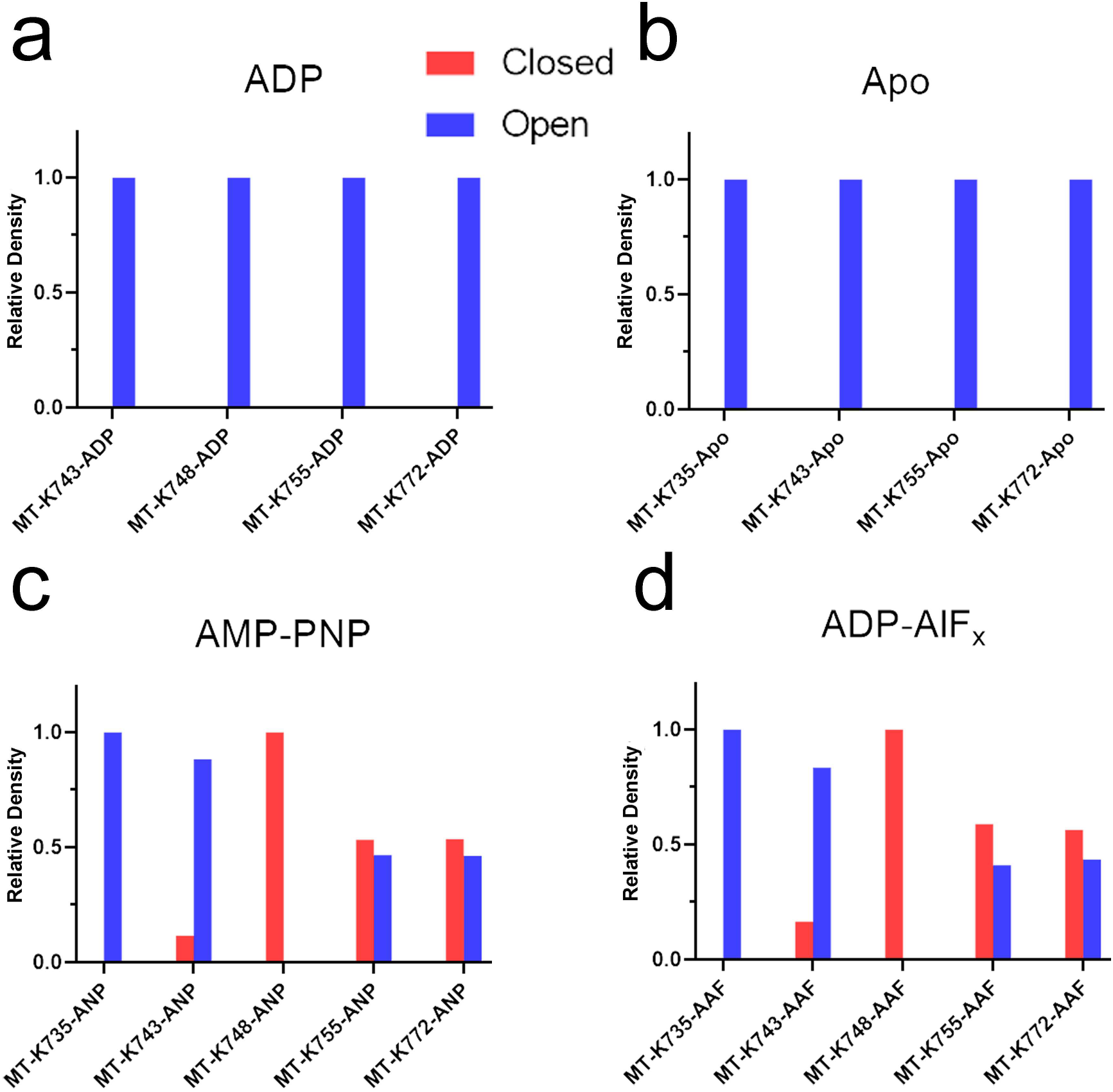
Cryo-EM map densities associated with open and closed conformations. Distinct and non-overlapping positions of KH0, KH1 and KH3 are found after aligning the structures to their corresponding β-tubulin subunit, one location corresponding to the open/open^*^ conformations and another corresponding to the closed conformation (Fig. 5f, g). **(a-d)** Relative density at the two alternate locations (closed or open/open*) for each cryo-EM map. Relative densities were calculated as RD_x_ = D_x_ / (D_O_ + D_C_), where Dx corresponds to the average density minus background at the open (D_o_) or closed regions (D_c_) (see methods). Note that in all the ADP and apo structures (a-b) the densities are associated with a single conformation (open). The MT-K748 associated densities are also associated with a single conformation, open or closed depending on nucleotide. On the other hand, the longer constructs K755 and K772 in the presence of AMP-PNP or ADP-AlF_x_ (c-d) present a near equal mix of open and closed conformation associated densities.

**Supplementary Fig. 5.**
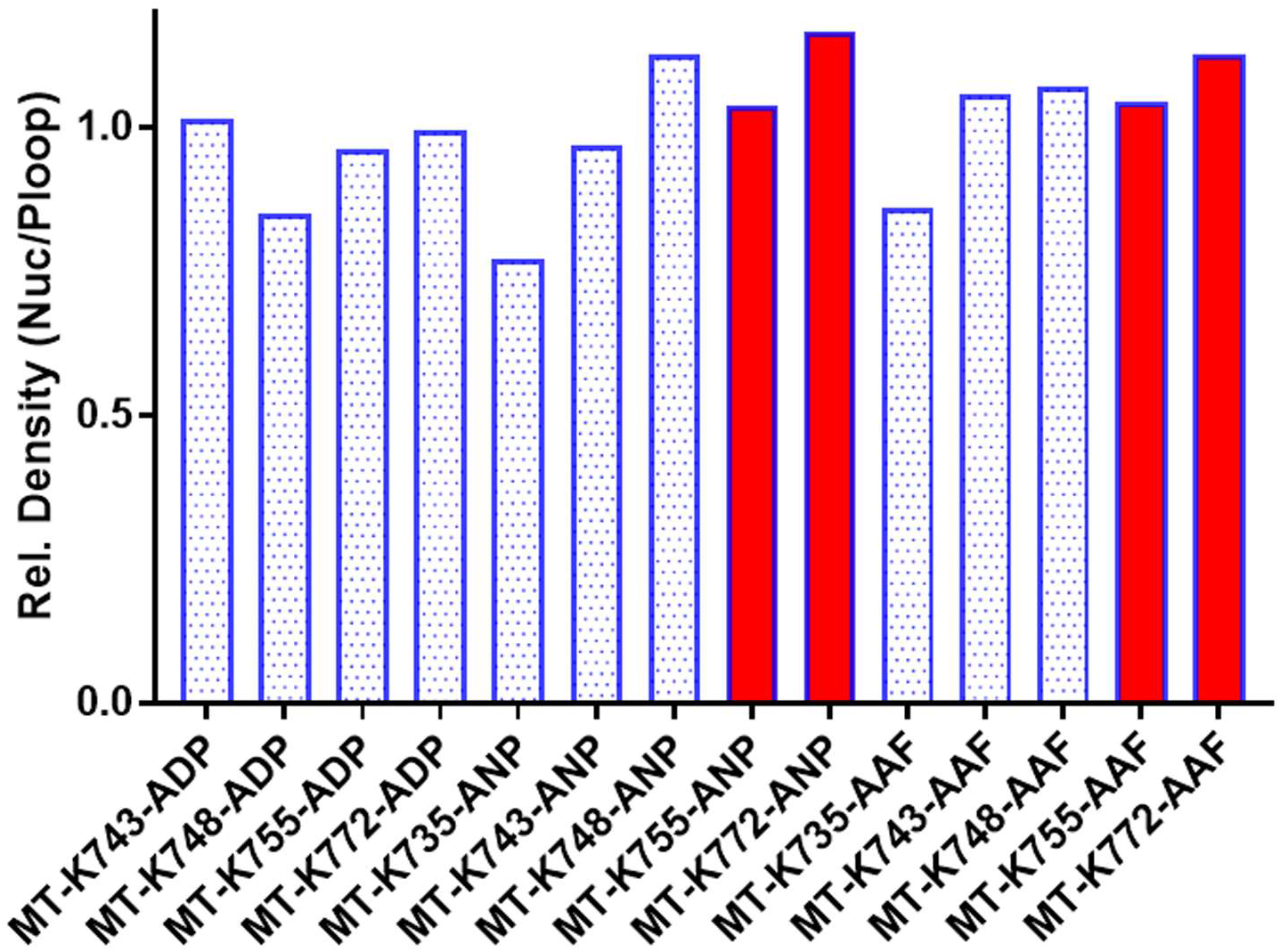
Relative density associated with the nucleotide (ADP or AMP-PNP) position of all cryo-EM maps. Relative nucleotide associated density (RDN) was calculated as RDN=DN/DPL where DN and DPL are the average density minus background at the nucleotide and P-loop locations (see methods). Note that the densities in the mixed open-closed conformations MT-K755-ANP/AAF and MT-K772-ANP/AAF structures (red bars) are similar or slightly higher than in the other structures.

**Supplementary Fig. 6.**
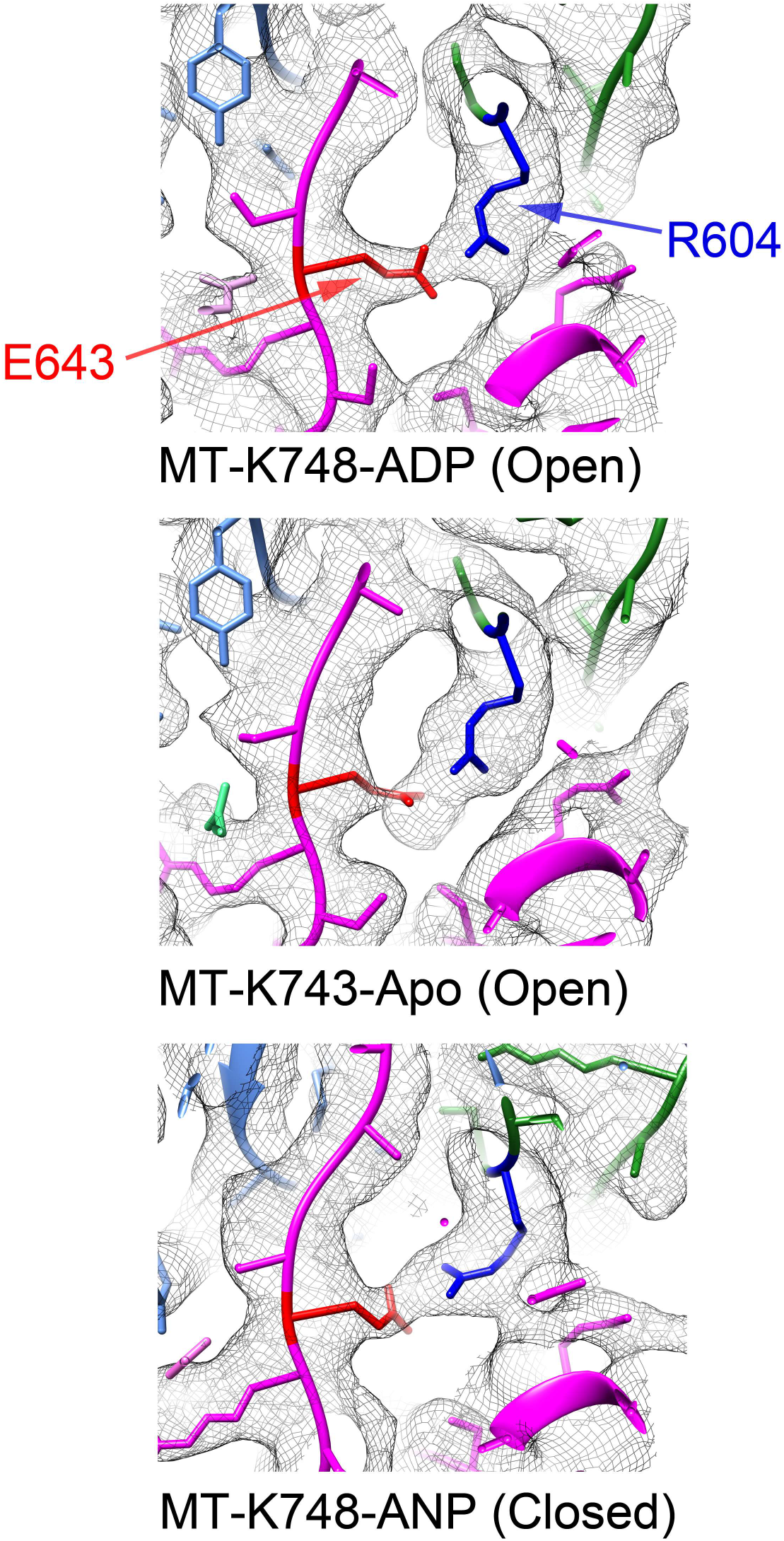
SW1-SW2 salt bridge. The salt bridge between residue R604 in SW1 and E608 is observed in all the MT-KIF14 complex structures regardless on whether the open or closed conformation (three examples shown). SW1 chain in green with R604 in blue, SW2 chain in magenta with E643 in red. Corresponding cryo-EM density is represented as an iso-density surface grey mesh.

**Supplementary Fig. 7.**
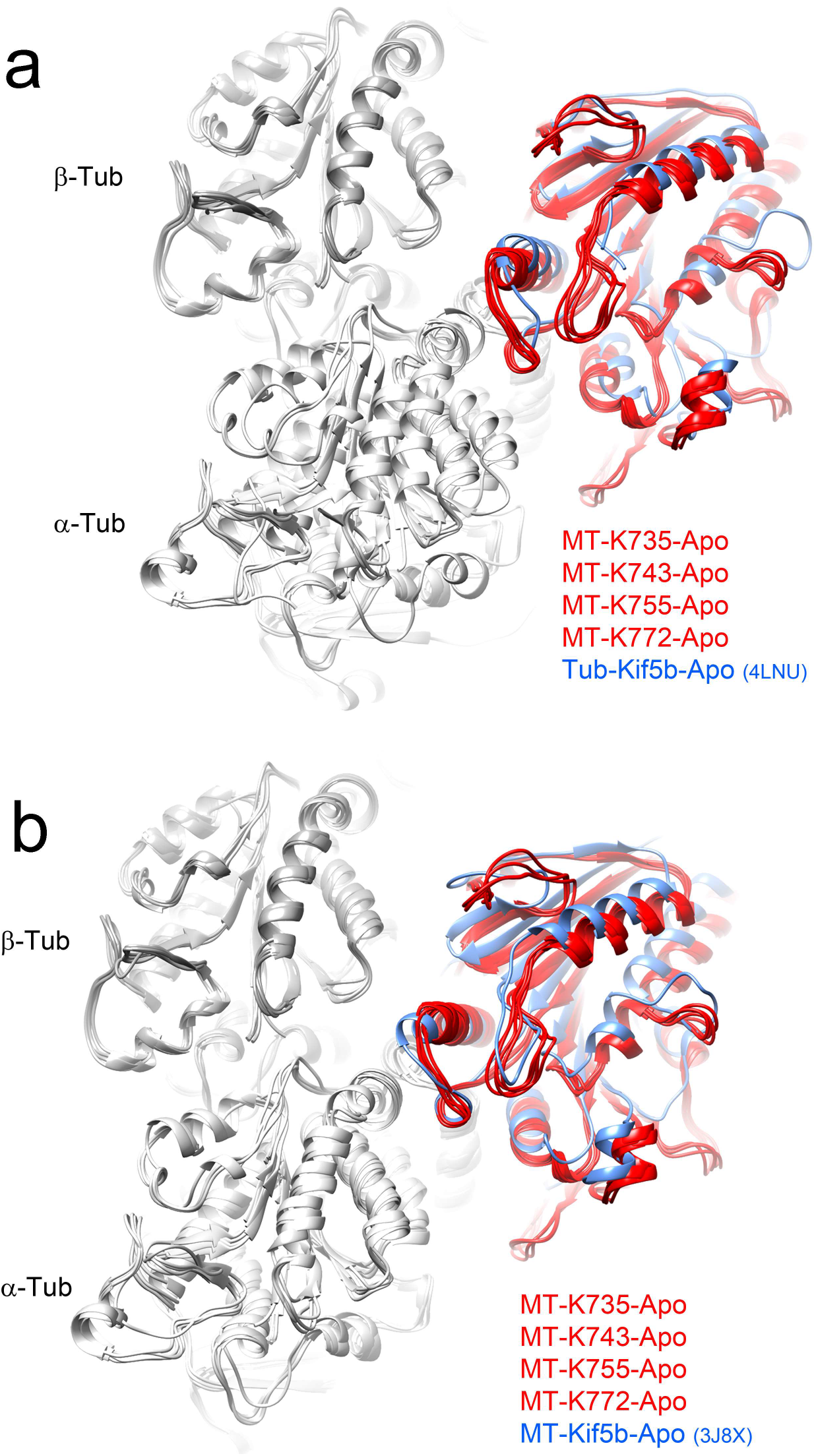
Microtubule-KIF14 and Tubulin-Kinesin-1 complex comparison. **(a)** Four microtubule-KIF14 complexes in the apo state structures superimposed with the tubulin-kif5b-apo complex (PDB: 4LNU). **(b)** Four microtubule-KIF14 complexes in the apo state structures superimposed with the microtubule-Kif5b-apo complex (PDB: 3J8X). Structures were superimposed by aligning the corresponding β-tubulin chains in the complexes.

**Supplementary Movie 1. Major conformations of the KIF14-microtubule complex through the ATPase cycle**. Movie made from seven structural models of KIF14 monomeric constructs, one from the microtubule unbound KIF14-ADP structure (4OZQ) and six from the microtubule-bound complexes (MT-748-ADP, MT-743-apo, MT-735-ANP, MT-748-ANP, MT-748-AAF, MT-735-AAF, MT-748-ADP). Structural transitions between states estimated by linear interpolation. Four successive different angles of view are displayed. The models were aligned on their β-tubulin subunit or the interaction area with β-tubulin in the case of the microtubule unbound KIF14 structure. β- and α-tubulin are colored in dark and light gray respectively. The KIF14 motor is colored according to the particular conformational change of the structure, semi-open in green, open in cyan, open* in blue and closed in red. Color saturation adjusted according to the amount of displacement between the open and closed structures (larger displacement more saturation). The side chains of key KIF14 nucleotide pocket residues, ARG 401, PRO 402, TYR 490, SER 489, SER 603 and ASN 599, are displayed as yellow sticks. The coordinated magnesium ion is displayed as a yellow sphere. The nucleotide is displayed in orange as sticks and the phosphate groups as spheres. The neck linker (only visible in the closed structures) is colored in red. Parts of KIF14 loop 8 (residues 543 to 551) and loop-10 (residues 618 to 627) are not displayed.

**Supplementary Movie 2. Kinesin dimer coordinated mechano-chemical cycle model.** Movie made from structural models of the MT-K755-ANP and MT-K755-AAF two-head bound states, the MT-K755-ADP and MT-K755-Apo one-head bound states and the microtubule unbound KIF14-ADP structure (4OZQ). Two successive angles of view are displayed. Structure representation and color scheme is as in Supplementary Movie 1. In the docked configuration the neck linker and coiled coil are colored in red while in the undocked configuration they are colored in pink. The neck-linkers and coiled coil of two proposed transient intermediates (trailing-open*-AAF with leading-open*-ANP and trailing-open-ADP with leading-open*-ANP), during and preceding detachment of the trailing head, are displayed in semitransparent grey. The relative time between states is only illustrative.

## Material and Methods

### Protein purification

KIF14 constructs were generated as GST-fusion proteins by amplification of the desired constructs from a plasmid containing the corresponding cDNA using Polymerase Chain Reactions (PCR). The plasmids were transformed into BL21 pLys Escherichia coli cells and were induced with 0.5 mM IPTG at 18°C overnight. GST-fusion proteins were purified on glutathione resins (Genscript) as previously described (Arora, Talje et al., JMB 2014). KIF14 constructs were cleaved on beads with PreScission protease [in cleavage buffer: 1X BRB80, 1 mM MgCl_2_, 100 mM KCl, 1 mM DTT, and 0.1 mM ATP] to generate the untagged motors. KIF14 constructs were flash frozen in small aliquots in liquid nitrogen, and stored in −80°C freezer.

Fifteen-protofilaments-enriched microtubules were prepared from porcine brain tubulin (Cytoskeleton, Inc. CO) as previously described (Benoit et al., 2018).

### ATPase activity assay

A malachite green-based phosphate detection assay was used to measure KIF14-mediated ATPase activity, as previously described (Talje et al. FEBS Lett. 2014, Arora et al., JMB 2014). Briefly, reactions were assembled in in BRB40-based depolymerization buffer (40 mM PIPES pH 6.8, 1 mM MgCl_2_, 1 mM EGTA, 20 μM taxol, 25 mM KCl, 0.25 mg/mL BSA, 1 mM DTT, 0.02% Tween), supplemented with 1 mM ATP, with 2 μM of taxol-stabilized MTs, and 50 nM KIF14 protein constructs. Basal activity of the KIF14 constructs was determined using the same reaction condition with no MTs added. Reactions were allowed to proceed for 10 min, quenched with perchloric acid and malachite green reagent (Arora et al., JMB 2014). The signal was quantified by the absorbance at 620 nm in a Genios Plus plate reader (Tecan).

### Cryo-EM KIF14-MT complexes sample preparation

K735-ANP, K735 apo and K772-ANP datasets were collected on untreated carbon/copper grids (Quantifoil R2/2 300 mesh). The grids used for the other 15 datasets were gold grids (UltrAuFoil R2/2 200 mesh), plasma cleaned just before use (Gatan Solarus plasma cleaner, at 15 Watts for 6 s in a 75% argon/25% oxygen atmosphere). Fresh microtubule preparation was performed on the day of the cryo-EM sample preparation and kinesin aliquots were thawed on ice just before use. All nucleotides stock solutions used were prepared with an equimolar equivalent of MgCl_2_ to ADP or AMP-PNP. Four μl of a microtubule solution at 2 to 5 μM tubulin (to account for the variability of microtubule binding on the grids) in BRB80 plus 20 μM paclitaxel) were layered onto the EM grid and incubated 1 min at room temperature. This microtubule solution also contains either AMP-PNP at 4 mM, ADP at 4 mM or ADP at 4 mM plus 2 mM AlCl_3_ and 10 mM KF. During the incubation time, a fraction of the thawed kinesin aliquot was diluted to prepare a 20 μM kinesin solution containing 20 μM Taxol and either of the four nucleotide conditions to be probed: 1) 4 mM ADP, 2) Apyrase: 5 × 10^−3^ units per μl (apo conditions), 3) 4 mM AMP-PNP, 4) 4 mM ADP plus 2 mM AlCl_3_, and 10 mM KF (ADP-AlF_x_ conditions). Then the excess microtubule solution was removed from the grid using a Whatman #1 paper. Four μl of the kinesin solution was then applied onto the EM grid, transferred to the chamber of a Vitrobot apparatus (FEI-ThermoFisher MA) at 100 % humidity were it was incubated for 1 min at room temperature, and blotted for 2.5 to 3 s with a Whatman #1 and −2 mm offset before plunge-freezing into liquid ethane. Grids were clipped and stored in liquid nitrogen until imaging in a cryo-electron microscope.

### Cryo-EM data collection

Data were collected at 300 kV on 3 Titan Krios microscopes equipped with K2 summit detectors (Supplementary Table 1). Acquisition was controlled using Leginon (Suloway et al., 2005) with the image-shift protocol and partial correction for coma induced by beam tilt (Glaeser et al., 2011). Data collection was mainly performed semi-automatically using 3 to 5 exposures per 2 μm diameter holes. The exposures were fractionated on 35 to 50 movie frames. The defocus ranges and cumulated dose are given in Table 1.

### Helical-single-particle 3D reconstruction

Movie frames were aligned with motioncor2 v1.0 or v1.1 generating both dose-weighted and non-dose weighted sums. All the datasets collected with a pixel size below 1 Å were corrected for magnification anisotropy (Supplementary Table 1). Before each of the corresponding session, a series of ~ 20 micrographs on a cross-grating calibration grid with gold crystals was used to estimate the current magnification anisotropy of the microscope using the program mag_distortion_estimate v1.0.1 (Grant and Grigorieff, 2015). Magnification anisotropy correction was performed within motioncor2 using the obtained distortion estimates (Supplementary Table 1). CTF parameters per micrographs were estimated with Gctf (v1.06) (Zhang, 2016) on aligned and non-dose-weighted movies average.

Images of 15R microtubules were processed using a helical-single-particle 3D analysis workflow as described (Benoit et al., 2018) with the following additions from step 5. At the end of step 5, the 2 independent refinements of half datasets are performed in Frealign (Grigorieff, 2016) limiting the refinement data to (1/8) Å^−1^ and run until no further improvement in resolution was detected. Number of particle images and asymmetric units included in the 3D reconstructions are reported in Table 1 and the particle-boxes sizes in Supplementary Table 1. A cleaning step was performed for several datasets (Supplementary Table 1) although the impact on resolution was negligible: particles for which the Euler angles are not compatible with 15R symmetry (either because not 15R or because they may be 15R particles poorly aligned) were discarded (< 4% of the data). If the resolution at that stage was at ~ 3.6 Å or better, per-particle image CTF values were refined. The datasets that underwent a CTF refinement step are listed in Supplementary Table 1. For such datasets, the defocus of each particles in each half-dataset were refined using one cycle of FrealignX (Grant et al., 2018) against their respective half-reconstruction and without particle alignment (defocus search parameters: step: 50 Å, range: 1250 Å). Because the defocuses of adjacent particles on a microtubule should vary smoothly, a moving median over 5 particles 82 nm apart was applied on each microtubule to assign their final per particle defocus values. After CTF refinement, further frealign cycles with limiting the refinement data to (1/5) Å^−1^ were run until no further improvement in resolution was detected. A 3D classification with Frealign in 2 classes to was used to improve the signal to noise (S/N) of the 748-ANP dataset. Eventually, whether a CTF refinement was performed or not, unmasked-unfiltered-unsharpened frealign reconstructions were obtained for each half dataset. Helical symmetry was then imposed in real space to each half map with relion_helical_toolbox.

To obtain a final locally filtered and locally sharpened map, the postprocessing of the pair of unfiltered and unsharpened half maps was performed in 2 main steps. (1) One of the 2 unfiltered half-map was low pass filtered to 15 Å and the minimal threshold value that does not show noise around the helical segment was used to generate a mask with relion-mask-create on 85 % of the helical segment on its helical axis (low pass filtration: 15 Å, extension: 10 pixels, soft edge: 10 pixels). This generous soft mask was used in blocres to apply a first local filtration of the map on 12 pixel size boxes (a purposely small box at stage to avoid over low pass-filtering). The unfiltered half maps were merged and corrected for the MTF of the detector by relion_postprocess without filtrations. The local resolution estimates by blocfilt were then applied to this merged map by locally low-pass filtering it with the program blocfilt. (2) The resulting map was then cropped in all dimensions with e2proc3d.py by ~27 % (to 484 pixel for 664 pixel sized boxes and to 382 pixels for 512 pixels sized boxes), keeping the helical segment centered. These cropped maps were used for local low-pass filtration and local sharpening in localdeblur (Ramírez-Aportela et al., 2019) with resolution search up to 25 Å. The localdeblur program converged with an optimum filtration for the tubulin which is too much sharpening for the kinesin, so maps of every localdeblur cycle were written to disk and the map with best filtration for the kinesin area was selected in the end. Eventually, the locally low-pass filtered and locally sharpened cropped map was inserted back into the full box (512 or 664 pixels sided boxes) and helical symmetry – not taken into account in the localdeblur filtering −was imposed.

### Cryo-EM resolution estimation

The final resolutions for each cryo-EM reconstruction were estimated from FSC curves generated with Relion 3.1 post-process program (FSC_0.143_ criteria, Table 1 and Supplementary Fig. 1). To estimate the overall resolution, these curves were computed from the two independently refined half maps (gold standard) using soft masks that isolate a single asymmetric unit containing a kinesin and a tubulin dimer. For each dataset using the PDB model obtained at the previous step, such a mask was generated with Eman1 pdb2mrc and Relion 3.1 relion_mask_create applied on the correctly positioned pdb2mrc output (low pass filtration: 15 Å, threshold: 0.1, extension: 2 pixels, soft edge: 6 pixels). FSC curves for the tubulin or kinesin parts of the maps were generated similarly using the corresponding subset of the PDB model to mask only a kinesin or a tubulin dimer (Table 1).

The atomic models, the cryo-EM final maps, the half maps, the mask isolating the single asymmetric units and the FSC curves were all deposited in the PDB and EMDB databases.

### Model building

Atomic models of the cryo-EM density maps were built as follow. First, atomic models for each protein chains were generated from their amino-acid sequence by homology modeling using Modeller (Fiser and Šali, 2003). Second, the protein chains were manually placed into the cryo-EM densities and fitted as rigid bodies using UCSF-Chimera (Pettersen et al., 2004). Third, the models were flexible fitted into the density maps using Rosetta for cryo-EM relax protocols (DiMaio et al., 2009; Wang et al., 2015) and the models with the best scores (best match to the cryo-EM density and best molprobity scores) were selected. Fourth, the Rosetta-refined models were further refined against the cryo-EM density maps using Phenix real space refinement tools (Adams et al., 2010). Fifth, the Phenix refined models were edited using Coot (Emsley and Cowtan, 2004). Several iterations of Phenix real space refinement and Coot editing were performed to reach the final atomic models.

For the K755 and K772 constructs in the AMP-PNP and ADP-AlF_x_, states the kinesin densities of each map were fitted with two coexisting ‘open’ and ‘closed’ structures. First the models corresponding to the MT-K743-ANP and MT-K748-ANP were rigidly fitted into the MT-K755-ANP and MT-K772-ANP maps and the MT-K743-AAF and MT-K748-AAF models into the MT-K755-AAF and MT-K772-AAF maps. The electron densities of the undocked neck-linker of the leading motor domain where built with Coot and the coiled-coil was modeled with CCBuilder 2.0 (Wood and Woolfson, 2018). Further refinement and editing of the structures was performed in Coot.

Atomic models and cryo-EM map figures were prepared with UCSF-Chimera (Pettersen et al., 2004) or VMD (Humphrey et al., 1996). Movies were made with VMD (Humphrey et al., 1996) and R (Ihaka and Gentleman, 1996).

### Cryo-EM density quantification

The relative intensity of different regions of the cryo-EM maps (Supplementary Figs. 3–5) were determined from density maps and the corresponding fitted atomic models using the <measure mapValues> command of UCSF-Chimera (Pettersen et al., 2004). When comparing the density of equivalent regions in different maps and atomic models the maps models were first aligned to their corresponding β-tubulin subunits (UCSF-Chimera matchmaker and matrixcopy commands). Densities were calculated as the average density near the atoms of the ligands or the backbone of specific KIF14 residues in the atomic models. Background density was estimated with a set of atoms placed near the selected residues but outside the area occupied by the atomic models. The KH0 region (Supplementary Fig. 3) was defined as residues 403-412 and the near mid and far locations respectively as the position of these residues in the MT-K748-ANP, MT-K735-ANP and MT-K748-ADP models. KH0-relative-density was estimated for all the maps as the intensity minus background at the near, mid and far locations over the total (near+mid+far). The Open and Closed conformation regions (Supplementary Fig. 4) were defined as residues 403-412, 501-515, 463-471 and 577-579 in the MT-K748-ADP and MT-K748-ANP models respectively. Relative density for each map was calculated as the average density at each position minus background (open or closed) over the total (open+closed). Nucleotide relative density (Supplementary Fig. 5) were calculated as the average density of the nucleotide atoms (ADP or AMP-PNP) over the average density at residues of the P-loop (KIF14 residues 476-494) in the same map.

